# NMR measurements of transient low-populated tautomeric and anionic Watson-Crick-like G·T/U in RNA:DNA hybrids: Implications for the fidelity of transcription and CRISPR/Cas9 gene editing

**DOI:** 10.1101/2023.08.24.554670

**Authors:** Or Szekely, Atul Kaushik Rangadurai, Stephanie Gu, Akanksha Manghrani, Serafima Guseva, Hashim M. Al-Hashimi

## Abstract

Many biochemical processes use the Watson-Crick geometry to distinguish correct from incorrect base pairing. However, on rare occasions, mismatches such as G•T/U can transiently adopt Watson-Crick-like conformations through tautomerization or ionization of the bases, giving rise to replicative and translational errors. The propensities to form Watson-Crick-like mismatches in RNA:DNA hybrids remain unknown, making it unclear whether they can also contribute to errors during processes such as transcription and CRISPR/Cas editing. Here, using NMR *R*_1ρ_ experiments, we show that dG•rU and dT•rG mismatches in two RNA:DNA hybrids transiently form tautomeric (G^enol^•T/U ⇄G•T^enol^/U^enol^) and anionic (G•T^−^/U^−^) Watson-Crick-like conformations. The tautomerization dynamics were like those measured in A-RNA and B-DNA duplexes. However, anionic dG•rU^−^ formed with a ten-fold higher propensity relative to dT^−^•rG and dG•dT^−^ and this could be attributed to the lower *pK*_a_ (Δ*pK*_a_ ∼0.4-0.9) of U versus T. Our findings suggest plausible roles for Watson-Crick-like G•T/U mismatches in transcriptional errors and CRISPR/Cas9 off-target gene editing, uncover a crucial difference between the chemical dynamics of G•U versus G•T, and indicate that anionic Watson-Crick-like G•U^−^ could play a significant role evading Watson-Crick fidelity checkpoints in RNA:DNA hybrids and RNA duplexes.

## INTRODUCTION

Watson-Crick base pairs (bps) share a common conformation typically referred to as the ‘Watson-Crick geometry’(1) (Figure 1A). Because all other twelve mismatches adopt alternative conformations, many enzymes use the Watson-Crick geometry to differentiate between correct and incorrect pairings, thereby increasing the fidelity of processes such as replication, transcription, translation, and CRISPR/Cas9 genome-editing.(2-6) However, on rare occasions, some mismatches can transiently adopt a Watson-Crick-like conformation through tautomerization or ionization of the bases (Figure 1B).(1,7) Masquerading as Watson-Crick bps, these alternative conformations of mismatches can evade fidelity checkpoints, resulting in replicative,(8-10) transcriptional,(11-13) or translational(14,15) errors and potentially CRISPR off-target editing(16) (Figure 1C).

**Figure 1.**
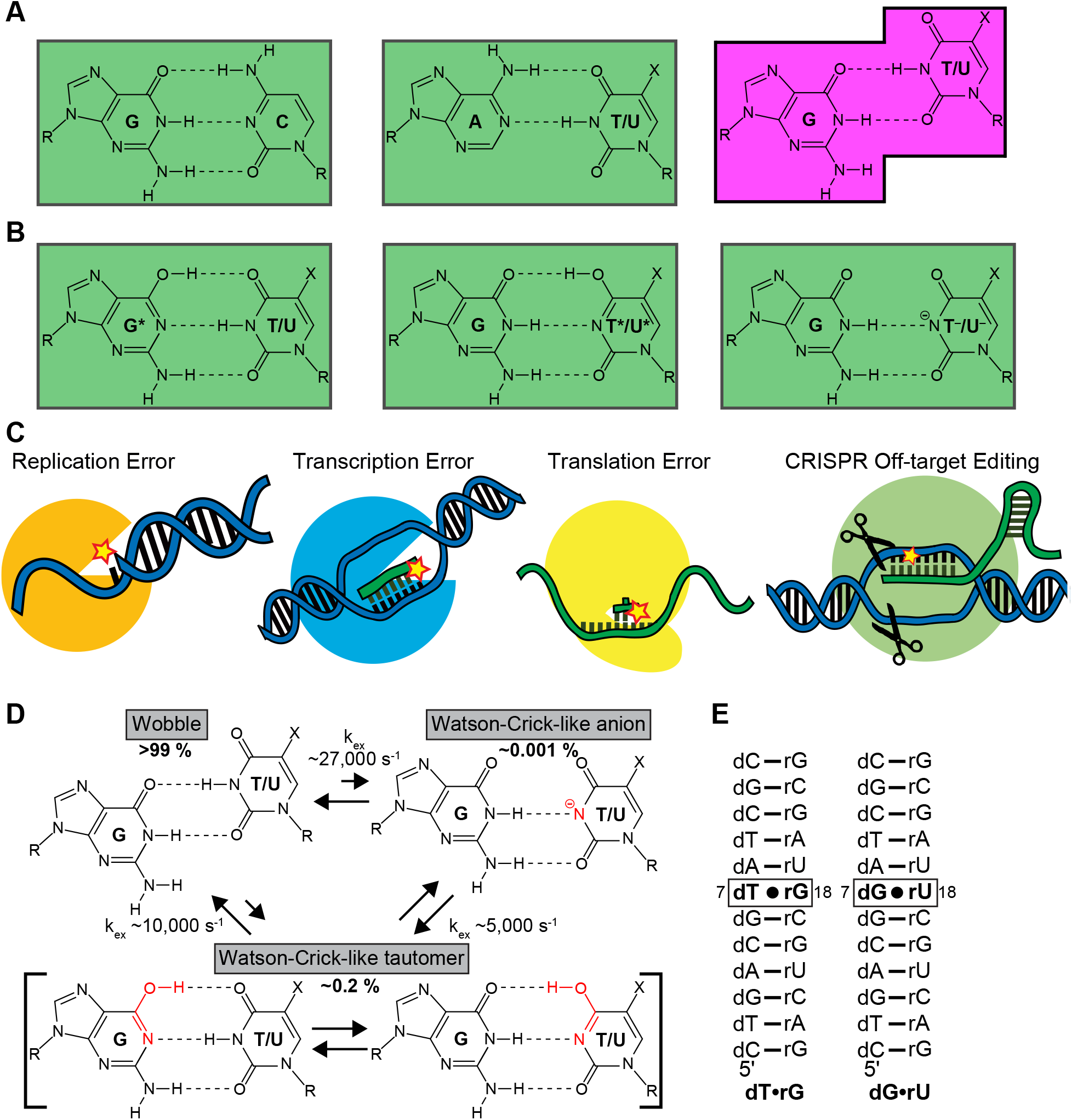
G•T/U mismatch conformational dynamics lead to the misincorporation of mismatches and errors during biochemical processes. (A) Canonical Watson-Crick base pairs form a common ‘Watson-Crick’ geometry (indicated by green rectangle) whereas mismatches such as G•T/U form alternative non-Watson-Crick conformations (magenta). X = methyl for T in DNA and hydrogen for U in RNA and R = sugar. (B) G•T/U can form Watson-Crick-like conformations through tautomerization or ionization of the bases. (C) By evading fidelity checkpoints, Watson-Crick-like G•T/U conformational states can contribute to replicative, transcriptional, and translational errors as well as CRISPR off-target editing. (D) Conformational exchange between wobble and Watson-Crick-like geometries of G•T/U mismatches formed by tautomerization (G•T^enol^/U^enol^ and G^enol^•T/U) and ionization (G•T/U^−^) of the bases as deduced using NMR *R*1ρ relaxation dispersion experiments in a prior study.(45) (D) The RNA:DNA hybrid duplexes used in the current study to examine the dynamics of dT•rG and dG•rU. Isotopically labeled nucleotides (^13^C/^15^N labeled dT and dG and ^15^N labeled rG and rU) are in bold.

Among twelve possible mismatches, G•T/U is particularly noteworthy as it is one of the most frequently misincorporated mismatches robustly across replication,(17-21) transcription,(22,23) and translation.(24-26) It is also one of the most commonly accommodated mismatches during CRISPR/Cas off-target editing.(16,27-31) Several studies implicate a role for Watson-Crick-like conformational states in these high-frequency errors (Figure 1D). Modifications stabilizing either the tautomeric or anionic Watson-Crick-like G•T/U also robustly increase the G•T/U misincorporation probability across replication,(8,32-34) translation,(35-37) and transcription.(11-13) Many crystal structures have been reported for DNA polymerase(10,38,39) and the ribosome(5,40-42) bound to G•T/U in catalytically active conformations, with the mismatch adopting a Watson-Crick-like conformation. These crystal structures could not distinguish between tautomeric and anionic species of the Watson-Crick like mismatch. Furthermore, they do not provide the kinetic information needed to assess the role of these states in determining the frequency of replicative and translational errors quantitatively. More recently, Nuclear Magnetic Resonance (NMR) studies (43,44) successfully resolved tautomeric and anionic Watson-Crick-like G•T/U bps forming transiently in low-abundance in B-DNA and A-RNA duplexes under solution conditions. The kinetic rates with which these Watson-Crick-like states formed could quantitatively explain the slower rate of G•T misincorporation relative to Watson-Crick bps and how the rate varies with pH and chemical modifications (Figure 1D).(45)

Transcriptional errors are also of great biological interest not only because they can be associated with dysregulated cellular functions such as splicing defects,(46,47) changes in cell metabolism,(48) and formation of aberrant proteins which misfold or aggregate,(49) leading to onset of neurodegenerative diseases,(50) but also because they can introduce variability into populations of cells, potentially offering a survival advantage in fluctuating environments.(51-53) However, in contrast to DNA and RNA, much less is known about the occurrence of Watson-Crick-like G•T/U conformational states in RNA:DNA hybrids and their potential roles in transcription fidelity. There are no crystal structures to date of RNA polymerase bound to G•T/U mismatches. In addition, NMR measurements of Watson-Crick-like G•T/U dynamics have not yet been performed on RNA:DNA hybrids. However, Watson-Crick-like rU•dG mismatches have recently been reported in two crystal structures of catalytically active CRISPR/Cas9-DNA ternary complex in the RNA:DNA hybrid within the Cas9 binding region formed by the single-guide RNA and its DNA target.(16) These structures suggest that Watson-Crick-like G•T/U can form in RNA:DNA hybrids and potentially contribute to undesirable off-target cleavage by CRISPR/Cas with important implications for therapeutic applications (Figure 1C).

The dynamics of tautomerization and ionization in the context of RNA:DNA hybrids is also interesting given the possibility for asymmetric dynamics when comparing dG•rU versus dT•rG,(54) which could in turn give rise to strand-specific errors. Here, we used off-resonance spin relaxation in the rotating frame (*R*_1ρ_) relaxation dispersion (RD) NMR experiments(55,56) to assess formation of Watson-Crick-like dG•rU and dT•rG bps in two RNA:DNA hybrids (Figure 1E). We find that dG•rU and dT•rG transiently form tautomeric conformations (G^enol^•T/U ⇄G•T^enol^/U^enol^) with dynamics like those measured in B-DNA and A-RNA duplexes. While dT•rG also formed anionic Watson-Crick-like dT^−^•rG with dynamics like those measured in B-DNA and A-RNA, anionic dG•rU^−^ formed with a ten-fold higher forward rate and population relative to dT^−^•rG. The high propensity to ionize dG•rU could be attributed to the lower *pK*_a_ (Δ*pK*_a_ ∼0.4-0.9) of U versus T. Our results suggest plausible roles for Watson-Crick-like G•T/U in transcriptional errors and CRISPR/Cas9 off-target gene editing, uncover a crucial difference between the chemical dynamics of G•U versus G•T, and indicate that anionic Watson-Crick-like G•U^−^ could play a significant role evading Watson-Crick fidelity checkpoints in RNA:DNA hybrids and RNA duplexes.

## MATERIALS AND METHODS

### Sample preparation

#### NMR Buffers

The buffer used for RD NMR experiments consisted of 15 mM sodium phosphate, 25 mM sodium chloride and 0.1 mM ethylenediaminetetraacetic acid (EDTA) in 90% H_2_O:10%D_2_O at pH 7.4 or at pH 7.8. The pH for buffers used in NMR titrations of NTPs was adjusted using sodium hydroxide (NaOH), while keeping the sodium ion concentration constant at 25 mM. All other components were as in the RD NMR buffers.

### Preparation of NMR samples

All the duplex RNA:DNA hybrid samples were prepared by mixing the two complementary strands in a 1:1 ratio at a concentration of ∼1 mM, heating to 95 °C for 5-10 min, followed by slow annealing at room temperature for ∼1 hr. NMR samples were buffer exchanged into NMR buffer (pH 7.4 or pH 7.8) using Amicon Ultra-15 centrifugal concentrators (3-kDa cutoff, Millipore Sigma) to a final nucleic acid concentration of ∼1 mM. Extinction coefficients used to measure oligonucleotide sample concentrations were estimated using the ADT Bio Oligo Calculator (https://www.atdbio.com/tools/oligo-calculator). Oligonucleotide sample concentrations ranged from 0.5 to 0.9 mM, and the sample pH was measured using a thin pH electrode.

### ^13^C/^15^N-labeled oligonucleotides

The DNA single-strands dCTGACG**T**ATCGC and dCTGACG**G**ATCGC were purchased from Yale Keck Oligonucleotide Synthesis Facility with cartridge purification in which residue dT7 or dG7 (in bold) were uniformly ^13^C/^15^N-labeled. The RNA single-strands rGCGAU**G**CGUCAG and rGCGAU**U**CGUCAG were synthesized in-house with a MerMade 6 Oligo Synthesizer in which residue rG18 or rU18 (in bold) were ^15^N site-specifically labeled. Standard RNA (n-ac-rA, n-ac-rG, n-ac-rC and rU, Chemgenes) and ^15^N site-specifically labeled RNA (^15^N-N1-rG and ^15^N-N3-rU) phosphoramidites,(57) and 1000 Å columns (Bioautomation) were used with a coupling time of 6 min, with the final 5′-dimethoxy trityl (DMT) group retained during synthesis. The oligonucleotides were cleaved from the supports (1 μmol) using ∼1 ml of AMA (1:1 ratio of ammonium hydroxide and methylamine) for 30 min and deprotected at room temperature for 2 h. The RNA single strands were dried under airflow to obtain oligonucleotide crystals. They were then dissolved in 115 μl DMSO, 60 μl TEA and 75 μl TEA.3HF and heated at 65°C for 2.5 h for 2′-O deprotection. The samples were then neutralized using 1.75 ml of Glen-Pak RNA quenching buffer, loaded onto Glen-Pak RNA cartridges for purification and were subsequently ethanol precipitated.

### Isotopically enriched NTP samples

Uniformly ^13^C/^15^N enriched rUTP and dTTP samples were purchased (Cambridge Isotope Laboratories) and dissolved using the NMR buffer (25 mM sodium chloride, 15 mM sodium phosphate, 0.1 mM EDTA and 10% D_2_O at variable pH values. The pH of the resulting sample was measured using a thin pH electrode: 6.92, 7.41, 8.0, 8.13, 8.15, 8.22, 8.24, 8.27, 8.42, 8.78, 8.87, 9.10, 9.20, 9.48, 9.93, 10.4, 11.2, 12.9, 13.5 (rUTP) and 6.89, 7.40, 8.08, 8.39, 8.62, 9.65, 9.81, 10.3, 10.8, 11.1, 11.5, 11.8, 12.0, 12.9, 13.6 (dTTP), and were adjusted using NaOH.

## NMR experiments

All NMR experiments were carried out at 25 °C (unless stated otherwise) on a Bruker Avance III 700 MHz spectrometer equipped with a HCN cryogenic probe.

### Resonance assignment

Assignments of imino resonances for dG•rU and dT•rG were easily obtained since they were site-specifically labeled (Supplementary Figure S1). The assignment for all other resonances was obtained using [^1^H, ^1^H] homonuclear Overhauser effect spectroscopy (NOESY) and aromatic [^13^C, ^1^H] heteronuclear correlation experiments. All the NMR spectra were processed using NMRpipe(58) and analyzed using SPARKY (T. D. Goddard and D. G. Kneller, SPARKY 3, University of California, San Francisco).

### ^13^C/^15^N R_1ρ_ relaxation dispersion

Off-resonance ^15^N *R*_1ρ_RD experiments(55,56) were implemented using a 1D selective excitation scheme as described in prior studies.(59,60) The spin-lock powers (ω_1_/2π) ranged from 400 to 2000 Hz, while the offsets ranged from ±3.5 times the spin-lock power, respectively (Supplementary Table S1). For each resonance 3-4 delay times were used during the relaxation period with maximum duration up to 120 ms.

### Analysis of R_1ρ_data

The *R*_1ρ_ data was analyzed as described previously.(43) Briefly, 1D peak intensities as a function of delay times extracted using NMRPipe,(58) were fitted to a mono-exponential decay to obtain the *R*_1ρ_ value for the different spin-lock power and offset combinations. The error in *R*_1ρ_ was estimated using a Monte Carlo procedure as described previously.(61) Exchange parameters of interests, such as the population of the excited states (ESs) (*p*_ES1_/*p*_ES2_), the exchange rates between ESs and the ground state (GS) (*k*_ex,GS:ES1_/*k*_ex,GS:ES2_/*k*_ex,ES1:ES2,_*k*_ex_ = *k*_f_ + *k*_b_, in which *k*_f_ and *k*_b_ are the forward and backward rate constants, respectively), the chemical shift difference between the ESs and the GS (Δω_ES1_/Δω_ES2_) and transverse (*R*_2_) and longitudinal (*R*_1_) relaxation rates were then extracted by fitting the *R*_1ρ_ data for a given nucleus to the Bloch-McConnell equations(62) describing a 2-state or 3-state exchange with a triangular topology (Figure 2 and Supplementary Figure S2). *R*_2,GS_=*R*_2,ES1_=*R*_2,ES2_=*R*_2_and *R*_1,GS_=*R*_1,ES1_=*R*_1,ES2_=*R*_1_ were assumed during fitting. Owing to the presence of an equilibration delay (5 ms) in the pulse sequence,(59,60) initial magnetization at the start of the Bloch-McConnell simulations was assumed to be equilibrated between the GS and ESs. Combined fits of the *R*_1ρ_ data for multiple nuclei were performed by sharing *p*_ES1_/*p*_ES2_ and *k*_ex,GS:ES1_/*k*_ex,GS:ES2_/*k*_ex,ES1:ES2_.

**Figure 2.**
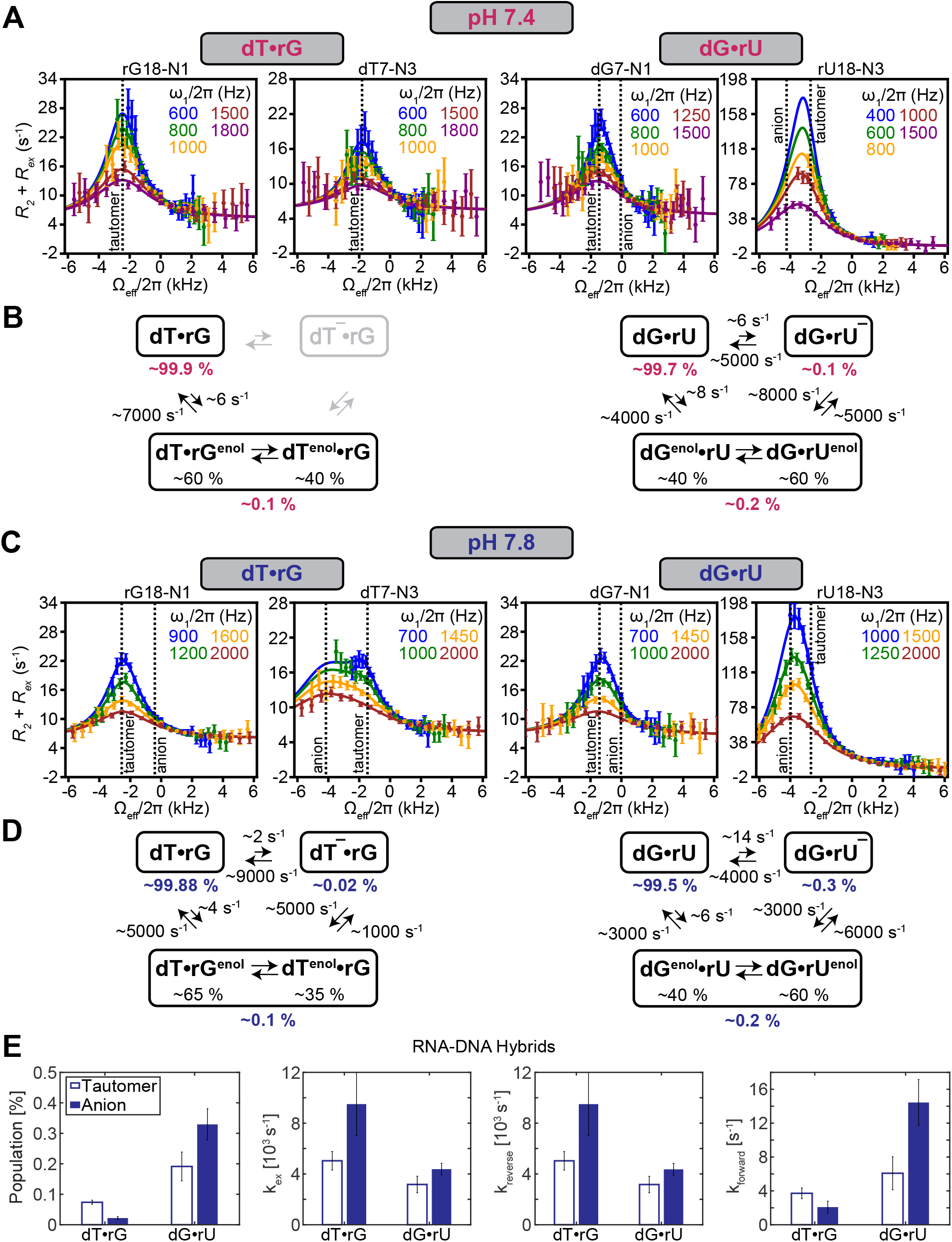
Measuring transient low-populated Watson-Crick-like G•T/U conformational states in RNA:DNA hybrids using NMR *R*1ρ relaxation dispersion. (A) Off-resonance ^15^N *R*_1ρ_ profiles for dT•rG and dG•rU measured at pH 7.4. Spin-lock powers are color-coded. Error bars represent the experimental uncertainty of the *R*_1ρ_ data and were obtained by propagating the error in *R*_1ρ_ as described in Materials and Methods. The solid lines denote fits of the data to a 2-state exchange model for dT•rG and a 3-state exchange model in a triangular topology for dG•rU. (B) The kinetic and thermodynamic parameters obtained from fitting the ^15^N *R*_1ρ_ profiles. (C, D) The corresponding (C) off-resonance *R*_1ρ_ profiles measured at pH 7.8 fit to a 3-state model with triangular topology along with the (D) derived kinetic and thermodynamic parameters. (E) Comparison of exchange parameters measured for the tautomeric (open bars) and anionic (full bars) conformational states in dT•rG and dG•rU at pH 7.8.

The initial alignment of magnetization during the Bloch-McConnell simulations was determined based on the *k*_ex_/Δω ratio of the majorly populated ES, as described previously.(61) The uncertainty in the exchange parameters was obtained using a Monte-Carlo scheme as described previously.(63) The fitted exchange parameters are summarized in Supplementary Tables S2 and S3.

Off-resonance *R*_1ρ_ profiles (Figure 2) were generated by plotting (*R*_2_ + *R*_ex_) = (*R*_1ρ_ -*R*_1_cos^2^ θ)/sin^2^ θ, where θ is the angle between the effective field of the observed resonance and the z-axis, as a function of Ω_eff_ = ω_obs_ -ω_RF_, in which ω_obs_ is the Larmor frequency of the observed resonance and ω_RF_ is the angular frequency of the applied spin-lock. Errors in (*R*_2_ + *R*_ex_) were determined by propagating the error in *R*_1ρ_ obtained as described above.

## *pK*_a_measurements and pH-dependent extrapolations

### Measurement of apparent pK_a_s for Watson-Crick-like G•T^−^/U^−^ using R_1ρ_

The population of the anionic Watson-Crick-like G•T^−^/U^−^ state deduced from fitting the *R*_1ρ_ data was utilized to calculate the apparent *pK*_a_ (*pK*_aapp_) using the Henderson-Hasselbalch equation (Eq. 1):

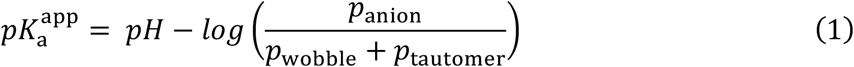

in which *p*_tautomer_ and *p*_anion_ are the equilibrium populations of the tautomeric and anionic ESs and *p*_wobble_ is the equilibrium population of the wobble GS. Similar *pK*_aapp_ values were obtained from measurements done for dG•rU at different pH conditions, validating the approach.

#### pH-dependent extrapolations

Prior RD data on B-DNA and A-RNA(45) (Supplementary Figure S3) as well as our current data on the dG•rU hybrid show that exchange kinetics between the GS and the neutral tautomeric ES1 species *k*_GS→ES1_ and *k*_ES1→GS_ are largely independent of pH (variations < 1.5 fold). Thus, *k*_GS→ES1_ and *k*_ES1→GS_ were assumed to be pH-independent.

The population of the anionic ES2 at any given pH was extrapolated using:

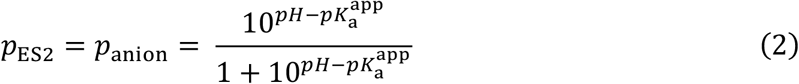

RD data also show (Supplementary Figure S3) that in the case of the anionic ES2, the backward rate *k*_ES2→ GS_ is largely pH independent (variations < 1.2 fold), consistent with a late transition state, and it is the forward rate *k*_GS→ES2_ that varies with pH giving rise to pH-dependent populations. By assuming the backward rate is independent of pH, we can deduce the forward rate for a given system at any pH based on the *pK*_a_ derived population. We implemented this model to extrapolate the pH dependent *k*_GS→ ES2_ as well as *k*_ES1→ ES2_ and *k*_ES2→ ES1_ describing minor exchange between ES1 and ES2 by first performing a linear fit of the rate constant versus pH based on RD data measured previously in B-DNA:(45)

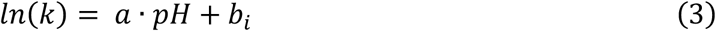

We then assumed that *a* is constant across *i* duplexes while *b*_i_ varies. Thus, the same pH-dependent fold change in the *k*_GS→ES2_, *k*_ES1→ES2_, and *k*_ES2→ES1_ measured in the reference DNA sample between two pH conditions were applied to the remaining *i* samples. Linear fits were done using Matlab®. It should be noted that the flux profiles through anion and tautomer as a function of *k*_2_ (Figure 4 and Supplementary Figure S7) obtained with this pH-dependent extrapolation were similar when instead using the *pK*_a_-derived *p*_ES2_ (Eq. 2) to determine the forward *k*_GS→ ES2_ rate 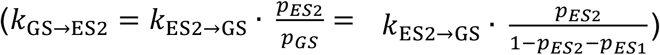 and do not affect any of the conclusions or claim made in our analysis.

##### Measurement of pK_a_s for [^13^C,^15^N]-rUTP and [^13^C,^15^N]-dTTP using NMR titrations

NMR titrations on the isolated nucleotides [^13^C,^15^N]-rUTP and [^13^C,^15^N]-dTTP were performed using the same buffer and temperature conditions as our NMR RD measurements by following the N3/C4 chemical shift changes with pH (Supplementary Figure S4). Peak positions (chemical shifts, ppm) were obtained using SPARKY (T. D. Goddard and D. G. Kneller, SPARKY 3, University of California, San Francisco), and inserted into Eq. 4 to get the normalized chemical shift perturbation, Δδ_obs_:(64)

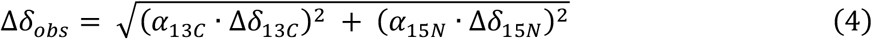

in which Δδ_13C_ / Δδ_15N_ are the ^13^C and ^15^N chemical shift difference relative to the N3/C4 resonance at pH 6.9, and α_13C_ / α_15N_ are the weight factors for ^13^C and ^15^N, respectively. These weight factors were estimated from the heteronuclear chemical shifts as observed for the 20 common amino acids in proteins using the BioMagResBank chemical shift database, where α_15N_ = 0.154 and α_13C=O_ = 0.341.(65)

To obtain the NTP *pK*_a_, the observed chemical shift difference was fitted to Eq. 5 using Matlab® non-linear fitting:

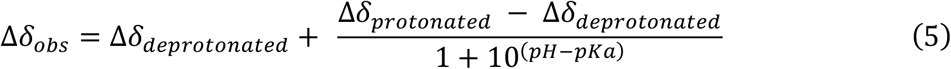

Which is a rearranged Henderson-Hasselbalch equation (Eq. 1). Δδ_deprotonated_ is the chemical shift difference of the deprotonated species (anion) and Δδ_protonated_ is the chemical shift difference of the protonated species. The populations in Eq. 1 can be expressed in terms of these chemical shift changes:(66)

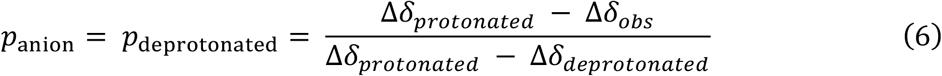

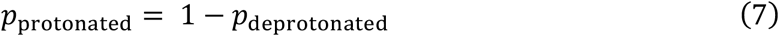

The chemical shift changes Δδ_obs_ and the fits to Eq. 5 are shown in Figure 3. The fitted *pK*_a_ values were 9.87 ± 0.01 and 10.32 ± 0.01 for rUTP and dTTP, respectively.

**Figure 3.**
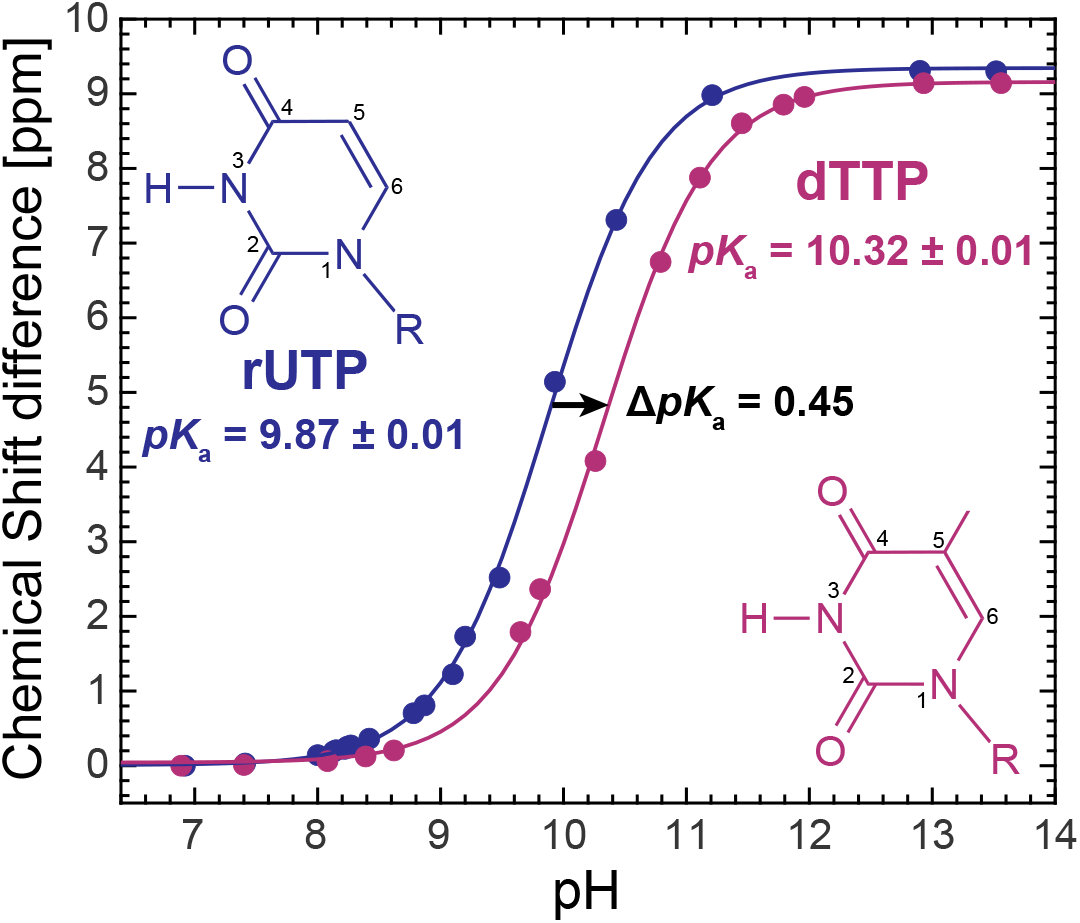
NMR titrations reveal a *pK*_a_ shift for dTTP relative to rUTP. NMR titrations of [^13^C,^15^N]-labeled rUTP (blue) and dTTP (purple), following the ^15^N and ^13^C chemical shifts of the N3/C4 resonance. The chemical structure of the NTP is indicated in the figure (R = triphosphate). Shown are the normalized changes in chemical shift with pH. Solid lines are fits to the Henderson-Hasselbalch equation, with *pK*_a_ = 9.87 ± 0.01 for rUTP, and *pK*_a_ = 10.32 ± 0.01 for dTTP. The fit error was estimated based on the square root of the diagonal values of the non-linear fitting covariance matrix. Experimental details and fitting equation are given in Materials and Methods.

**Figure 4.**
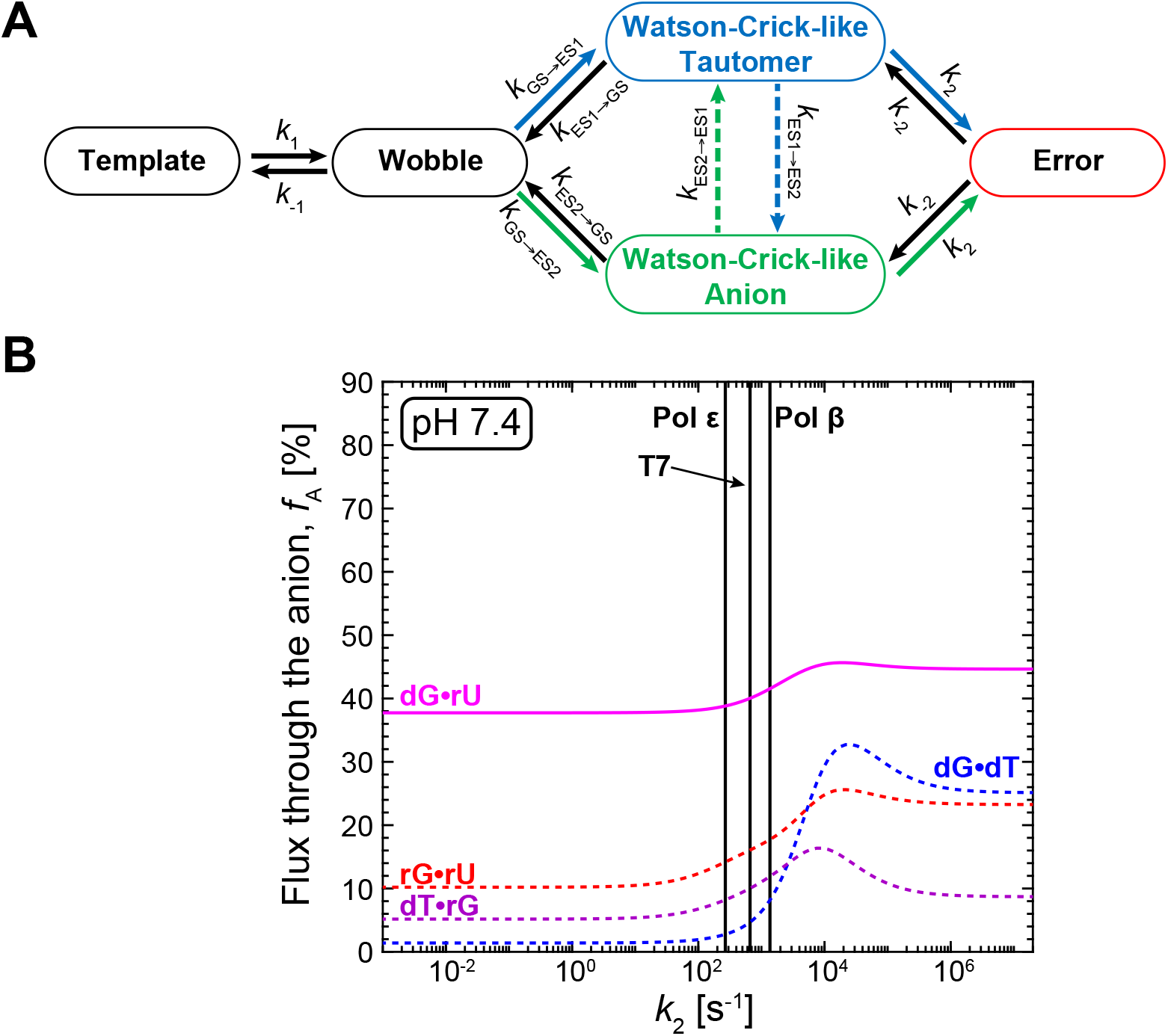
Anionic Watson-Crick-like G•U^−^ can significantly contribute to misincorporation errors. (A) Kinetic pathways for misincorporating the G•T/U mismatch via tautomeric (in blue) and anionic (in green) Watson-Crick-like G•T/U conformational states. Shown are the binding step (*k*_1_) to form the G•T/U wobble ground-state (GS); transition of the wobble into the tautomeric (*k*_GS-ES1_) and anionic (*k*_GS-ES2_) Watson-Crick-like G•T/U; and the kinetic step acting on the Watson-Crick-like species, which in DNA replication represents conformational changes in the DNA polymerase to form the catalytically active closed conformation. The binding step was assumed to be equal to that of DNA Polymerase ε.(45) Relative flux quantifying the fractional percentage (*f*_A_) of misincorporations proceeding through the anionic Watson-Crick-like G•U^−^ conformation as a function of varying *k*_2_ computed using the kinetic model presented in (A). Results are shown when using the chemical dynamics measured by NMR for dG•dT (DNA) and rG•rU (RNA) with a CGC trinucleotide sequence context(45) and dG•rU and dT•rG in the two RNA:DNA hybrids examined in this work at pH 7.4. The flux through the two pathways is computed using NMR-derived exchange rates (full lines), or extrapolated exchange rates (dotted lines, see Materials and Methods). The calculations assume that the Watson-Crick-like G•T/U is an obligatory intermediate during misincorporation. Vertical lines indicate *k*_2_ values for Pol ε,(68,69) T7,(70,71) and Pol β.(72)

### Equilibrium flux calculations

We used the kinetic scheme in Supplementary Figure S6A to compute the relative misincorporation flux through the tautomeric (*f*_T_) and anionic (*f*_A_) Watson-Crick-like conformational states.

The kinetic equations used to compute misincorporation flux:

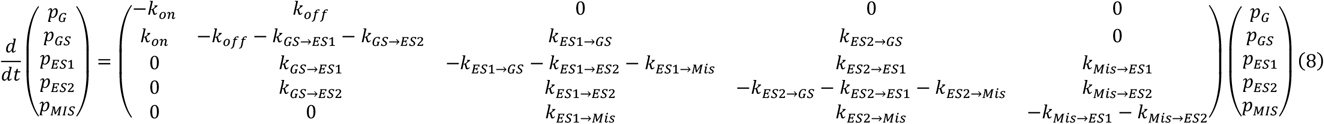

in which G represents the bound species before NTP binding; GS is the wobble G•T/U ground state; ES1 and ES2 are the tautomeric and anionic Watson-Crick-like G•T/U excited states respectively; and MIS is the misincorporated G•T/U for which the polymerase is in a catalytically active conformation. *k*_i→ j_ is the rate constant for transitioning between the *i* and *j* states and *p*_i_ is the population of the *i* state, and *k*_on_ and *k*_off_ are the binding on and off rates, respectively (*k*_1 and *k* -1_). The model assumes that misincorporation from the wobble GS is negligible; that the rate constants for misincorporating ES1 (*k*_ES1→MIS_ *k*_MIS→ES1_) and ES2 (*k*_ES2→MIS_; *k*_MIS → ES2_) are equal (*k*_ES1→MIS_ = *k*_ES2→ MIS_ ≡ *k*_2_; *k*_MIS→ES1_ = *k*_MIS→ ES2_ ≡ *k*_-2_); and that the tautomerization and ionization rates measured in the duplex by NMR approximate the rates in the polymerase active site.

Additionally, simulations show that the binding step has little to no effect on the relative flux through the anion species (*f*_A_, Supplementary Figure S6B). We therefore use a simplified model for the kinetic system:

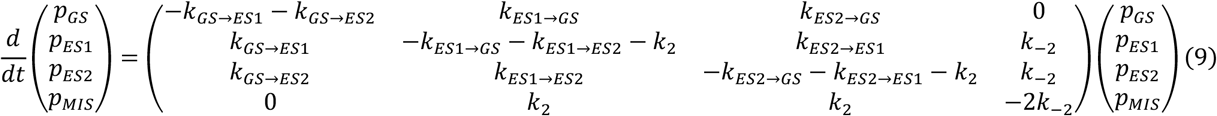

These differential equations were solved using an in-house Python 2.7 script utilizing Python ODE solver. The initial conditions were set so that the *p*_GS_ = 1 and *p*_ES1_ = *p*_ES2_ = *p*_MIS_ = 0, and the total simulation time was *t*_sim_ = 10 s, for which we assume the system has reached equilibrium, since *t*_sim_ >> *t*_eq_ (∼0.005 s for most systems) and so the extracted parameters are equilibrium populations. These computed equilibrium populations are then used below to compute the flux through each one of the ES pathways.

The net flux through parallel reaction paths (*F*_par_) is given by the sum over the flux through all available parallel reaction pathways:(67)

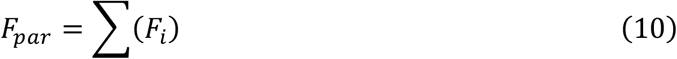

in which *F*_i_ is the flux through the pathway *i*. The flux of a serial reaction path (*F*_ser_), which is comprised of sequential reactions, is defined as:

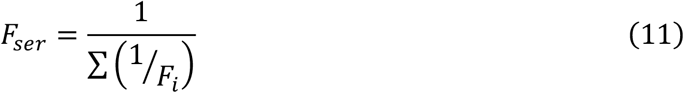

We treat the flux pathway through the tautomer (ES1) as a combination of two parallel pathways GS→ES1→MIS (pathway *F*_tt_) and GS→ES2→ES1→MIS (pathway *F*_ta_). Therefore, the tautomer flux is given by:

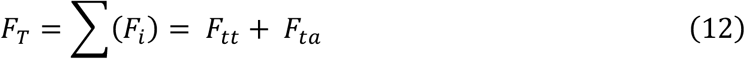

Likewise, we treat the flux pathway through the anion (ES2) as a combination of two parallel pathways GS→ES2→MIS (pathway *F*_aa_) and GS→ES1→ES2→MIS (pathway *F*_at_). The flux through the anion is then given by:

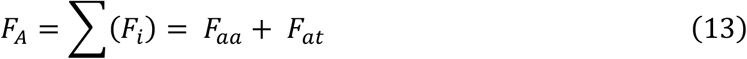

The relative flux through the anion and the tautomer is given by:

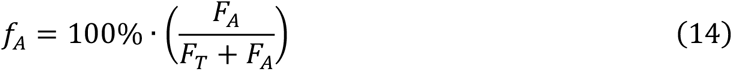

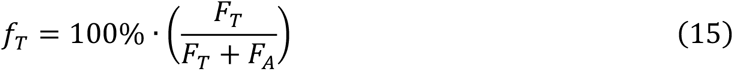

The flux through the different pathways *F*_tt_, *F*_ta_, *F*_aa_, and *F*_at_ are calculated as a combination of serial reaction pathways:

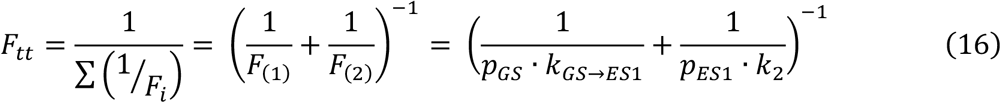

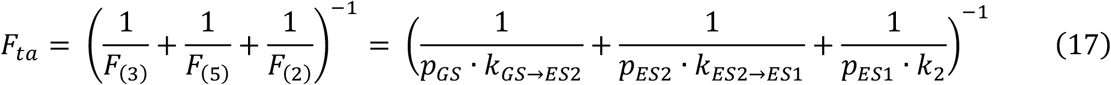

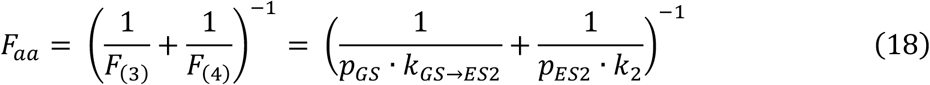

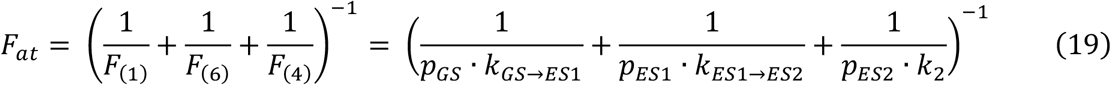

The values of *p*_GS_, *p*_ES1_ and *p*_ES2_ were computed by solving the kinetic equations in Eq. 9 at equilibrium (*t*_sim_ = 10 s); exchange rates *k*_GS→ ES1_, *k*_GS→ES2_, *k*_ES1→ES2_and *k*_ES2→ES1_ were obtained from fitting the NMR *R*_1ρ_ data; and *k*_2_ is polymerase-specific and was either taken from other kinetic studies, or varied as a free parameter.

This simplified model of misincorporation kinetics gives a good approximation for the relative contributions of the tautomeric and anionic states to misincorporation by DNA polymerase or by RNA polymerase. Similar results are obtained when calculating the flux through each pathway using the complete model (data not shown). Flux calculations were performed for misincorporation by the human DNAP Pol ε,(68,69) T7 DNAP(70,71) and rat Pol β,(72) for which the microscopic rate constants are known.

## RESULTS AND DISCUSSION

### dT•rG in an RNA:DNA hybrid forms tautomeric Watson-Crick-like conformations with dynamics similar to B-DNA and A-RNA

We used off-resonance spin relaxation in the rotating frame (*R*_1ρ_) NMR spectroscopy to study the dynamics dG•rU or dT•rG bps in two RNA:DNA hybrid duplexes previously used to measure the misincorporation probabilities of dG•rU and dT•rG during transcription.(54) In these NMR samples, residues forming the dG•rU and dT•rG bps were specifically ^13^C/^15^N labeled using chemical synthesis (see Materials and Methods). Based on the NMR chemical shifts and NOE-based distance connectivity (Supplementary Figure S1), dG•rU and dT•rG formed the expected wobble mismatches as the dominant conformation, with all other residues forming canonical Watson-Crick bps.

Prior *R*_1ρ_ NMR studies on B-DNA and A-RNA duplexes(43-45) showed that the wobble G•T/U ground-state (GS) exchanges with two short-lived low-abundance Watson-Crick-like ‘excited conformational states’ (ES); one representing two tautomeric states in rapid equilibrium on the NMR chemical shift timescale (G^enol^•T/U ⇄G•T^enol^/U^enol^), referred to as ‘ES1’, and one corresponding to an anionic species (G•T^−^/U^−^), referred to as ‘ES2’. Because these dynamics were observed robustly in B-DNA and A-RNA duplexes with comparable kinetic rates,(43-45) we expected they would also be observable in the RNA:DNA hybrids which typically adopt A-form-like conformations depending on sequence context.

We measured off-resonance ^15^N *R*_1ρ_ RD(55,56) for G-N1 and U/T-N3 (Figure 2A) under the same pH conditions (pH 7.4) used previously to measure misincorporation of G•T/U during transcription.(54) Based on prior dynamics measurements in B-DNA,(43-45) the population of the anionic Watson-Crick-like G•T^−^/U^−^ under these pH conditions should fall below the detection limits of the *R*_1ρ_ experiment, such that only the tautomeric Watson-Crick-like G•T/U is observable, simplifying analysis. The *R*_1ρ_ experiment measures the line-broadening contribution (*R*_ex_) to the transverse relaxation rate (*R*_2_) during a relaxation period in which a continuous radiofrequency (RF) field is applied with variable power (ω_1_) and frequency (ω_RF_). The RF field diminishes the *R*_ex_ contribution in a manner dependent on the magnitude of ω_1_ and ω_RF_ and the exchange parameters of interest. RD profiles are typically displayed by plotting *R*_2_ + *R*_ex_ versus ω_1_ and ω_RF_. For detectable exchange, a peak is typically observed centered at the difference between the chemical shift of the GS and ES (-Δω, assuming ω_GS_ = 0 and ω_ES_ = Δω). The dependence of *R*_2_ + *R*_ex_ (*R*_1ρ_) on ω_1_ and ω_RF_ can be fit to the Bloch-McConnell equations(73) describing n-site exchange to determine exchange parameters of interest.

Indeed, for both dT•rG and dG•rU, the U/T-N3 and G-N1 RD profiles (Figure 2A) indicated that the wobble GS exists in dynamic equilibrium with a short-lived lowly populated ES. The dT•rG RD profiles could be satisfactorily fit to a 2-state exchange model (Supplementary Figure S2A and Supplementary Table S2). The ES dG-N1 and dT-N3 chemical shifts (Δω = ω_ES_ – ω_GS_; Δω_G-N1_ = 34.9 ± 0.7 ppm and Δω_T-N3_ = 25.7 ± 0.8 ppm) deduced from the 2-state fit were as expected for two rapidly exchanging tautomeric species dT•rG^enol^ (∼60%) ⇄dT^enol^•rG (∼40%), as reported previously in B-DNA and A-RNA.(43,45) In addition, the population (*p*_ES_ = 0.083 ± 0.002 %) and exchange rate (*k*_ex_ = *k*_forward_ + *k*_reverse_ = 7,200 ± 500 s^-1^) deduced from the 2-state fit (Figure 2B) were similar to those measured previously for the tautomeric Watson-Crick-like dG•dT in B-DNA and rG•rU in A-RNA duplexes.(43,45) Thus, tautomeric Watson-Crick-like dT•rG can form in an RNA:DNA hybrid under solution conditions with kinetic and thermodynamic propensities similar to B-DNA and A-RNA duplexes.

### dG•rU in an RNA:DNA hybrid forms tautomeric Watson-Crick-like conformations and has a high propensity to form anionic dG•rU^−^

Unexpectedly, for the dG•rU mismatch in the second RNA:DNA hybrid, the dG-N1 and rU-N3 RD profiles called for a 3-state fit involving two ESs in a triangular kinetic topology (Supplementary Figure S2A, Supplementary Table S2). One ES had exchange parameters (Δω_G-N1_ = 21 ± 3 ppm, Δω_U-N3_ = 36 ± 4 ppm, p_ES_ = 0.19 ± 0.06 %, *k*_ex_ = 4,000 ± 900 s^-1^) consistent with two rapidly exchange tautomeric species; dG^enol^•rU (∼40%) ⇄dG•rU^enol^ (∼60%). Relative to dT•rG, the slightly higher population of dG•rU^enol^ relative to dG^enol^•rU measured in this hybrid is consistent with prior NMR studies(45) of dG•dT and rG•rU in B-DNA and A-RNA duplexes and can be attributed to the impact of the electron donating methyl group in dT, which destabilizes dT^enol^ relative to rU^enol^.(45,74)

The second ES had chemical shifts (Δω_G-N1_ = 1 ± 7 ppm and Δω_U-N3_ = 58 ± 7 ppm) consistent with an anionic Watson-Crick-like dG•rU^−^ with population *p*_ES_ = 0.12 ± 0.06 % and *k*_ex_ = 5,000 ± 2,000 s^-1^. The ^15^N chemical shifts as well as other lines of evidence(8,75,76) rule out an anionic inverted wobble conformation which has been reported in some crystal structures of DNA polymerase,(38,77) in complexes of the ribosome, mRNA and modified tRNA,(78,79) as well as in ribosomal RNA.(80) Furthermore, the anionic and tautomeric species exchange with *k*_ex,minor_ = 13,000 ± 6,000 s^-1^ (Figure 2B). Thus, dG•rU can form a tautomeric Watson-Crick-like conformation and also has a surprisingly high propensity to form anionic Watson-Crick-like dG•rU^−^ relative to both dT^−^•rG and dG•dT^−^.

We were able to confirm the higher propensity to form anionic dG•rU^−^ relative to dT^−^•rG by repeating the *R*_1ρ_ experiments at a slightly higher pH of 7.8 (Figure 2C, 2D, Supplementary Figure S2B, Supplementary Table S3). The anionic species was now observable in both dG•rU and dT•rG mismatches. The population of dG•rU^−^ increased ∼3-fold with increasing pH and was 10-fold higher (∼0.3 % vs. ∼0.02 %) than that of dT^−^•rG due to a 10-fold faster forward rate (∼14 s^-1^ vs. ∼2 s^-1^) (Figure 2E). The apparent *pK*_a_ (see Materials and Methods) for dG•rU was ∼10.3 compared to ∼11.5 for dT•rG. As expected, the population of the tautomeric species did not change significantly (<1.5-fold) with pH.

### The high propensity to form dG•rU^−^ is in part driven by the lower *pK*_a_ of rU-N3 relative to dT-N3

What accounts for the higher propensity to form Watson-Crick-like dG•rU^−^ relative to dT^−^•rG? Based on prior *pK*_a_ measurements(81,82) on rU, dU, rT and dT mono-phosphate nucleotides, the *pK*_a_ of U-N3 is robustly lower than T-N3 by 0.5 – 0.9 units. The lower *pK*_a_ has been attributed(83) to the absence of the electron donating methyl group. We confirmed these prior findings by measuring the *pK*_a_s for the nucleotides [^13^C,^15^N]-rUTP and [^13^C,^15^N]-dTTP, using N3/C4 NMR chemical shift titrations in the same buffer and temperature conditions used to measure the dG•rU and dT•rG dynamics (Figure 3, Supplementary Figure S4). Indeed, the *pK*_a_ measured for rUTP was ∼0.5 units lower than that of dTTP, corresponding to a ∼4-fold higher propensity for rUTP to deprotonate relative to dTTP.

If the higher propensity to form dG•rU^−^ relative to dT^−^•rG is due to the lower *pK*_a_ of rU-N3 versus dT-N3, we might expect that for the same sequence context, rG•rU in A-RNA would have a higher propensity to deprotonate and form rG•rU^−^ relative to dG•dT^−^ in B-DNA. While such a head-to-head comparison had never been made before, prior NMR studies did measure the population of this anionic species at the same pH conditions (pH 8.4) in B-DNA and A-RNA duplexes that share a common CGC trinucleotide sequence context. Indeed, reexamining this data we find that the rG•rU^−^ population was 10-fold higher than that of dG•dT^−^ (Supplementary Figure S5) and this was due to a 10-fold slower backward rate. Thus, while additional studies are needed to fully examine the contribution of sequence context, the higher propensity to form dG•rU^−^ relative to dT^−^•rG in the two RNA:DNA hybrids is most likely dominated by the intrinsically lower *pK*_a_ of U versus T.

### Anionic G•U^−^ can be a substantial driver of errors in RNA:DNA hybrids and RNA duplexes

We previously(45) developed a quantitative and predictive kinetic model for dG•dT misincorporation during replication by high fidelity DNA polymerases which involves Watson-Crick-like tautomeric and anionic species as obligatory intermediates. In this model (Figure 4A), following initial binding in a wobble conformation with DNA polymerase in the open state, dG•dTTP (or dT•dGTP) forms either the Watson-Crick-like tautomeric or anionic conformation. All subsequent steps, including conformational changes in DNA polymerase (*k*_2_) to form the closed catalytically active state and the phosphodiester bond formation (*k*_3_) proceed at kinetic rates equal to those measured for canonical Watson-Crick bps. Based on this model, we previously estimated that at physiological pH∼7, dG•dT misincorporation during DNA replication proceeds primarily (>90%) through the tautomeric Watson-Crick-like G•T.(45)

Given its comparatively higher propensity to form in RNA and RNA:DNA hybrids, anionic Watson-Crick-like G•U^−^ could have a more substantial role driving translational and transcriptional errors as well as CRISPR/Cas off-target editing relative to DNA replication. We used our kinetic model for DNA misincorporation as a basis for estimating the magnitude of this error contribution. To generalize our findings, we performed simulations varying the kinetic rate of the step (*k*_2_) acting on the Watson-Crick-like conformation. Kinetic simulations show that other steps including binding have little consequence on the relative flux through the anionic versus tautomeric species (Supplementary Figure S6). Thus, the conclusions made can be generalized to a wide variety of biochemical processes employing different mechanisms so long as the Watson-Crick-like state is an obligatory intermediate.

Shown in Figure 4B is the anionic error contribution defined as the relative flux during misincorporation through the anion, *f*_A_, as a function of varying *k*_2_ for the two hybrid duplexes dG•rU and dT•rG at physiological pH. The flux represents the fractional percentage of misincorporations proceeding via the anionic Watson-Crick-like species. For comparison, data is also shown for dG•dT in B-DNA and rG•rU in A-RNA with the common CGC trinucleotide sequence context. For the systems in which the anionic conformation was not directly detected by NMR at pH 7.4 (hybrid dT•rG, RNA rG•rU, and DNA dG•dT), kinetic rates were extrapolated based on pH-dependent data(45) (see Materials and Methods and Supplementary Figure S3 for details).

In general, the dependence of flux on *k*_2_ varies between two limits. In the ‘thermodynamic limit’, *k*_2_ << *k*_ES2→GS_ and *k*_ES1→GS_, and the step acting on the Watson-Crick-like state is rate limiting. Under this limit, the wobble-to-Watson-Crick-like dynamics attain thermodynamic equilibrium, and the flux through the anionic species is approximately given by the fractional population of the anionic species relative to both the anionic and tautomeric conformation i.e. *f*_A_ ∼ *p*_ES2_/(*p*_ES1_ + *p*_ES2_).(45) In the ‘kinetic limit’, *k*_2_ >> *k*_ES2→GS_ and *k*_ES1→GS_, and formation of the Watson-Crick-like conformation is rate limiting. Under this limit, the flux through the anionic species is approximately given by the relative magnitude of the forward rates for the two species i.e. *f*A ∼ *k*GS→ES2/(*k*GS→ES1 + *k*GS→ES2) (assuming no minor exchange). Also shown on Figure 4B are the *k*2 values for DNA polymerase ε,(68,69) T7 DNA polymerase,(70,71) and rat DNA polymerase β (72).

Strikingly, we find that the error contribution due to anionic dG•rU^−^ in the RNA:DNA hybrid at pH 7.4 was around *f*A ∼40% robustly across different *k*2 values. In contrast, the flux for dT•rG in the second hybrid was *f*A ∼10% and for dG•dT it was <8% for *k*2 values corresponding to DNA polymerases. The flux for rG•rU in RNA was also comparatively high with *f*A ∼14-18%. These trends, including the greater flux through anionic dG•rU^−^ in the RNA:DNA was observed robustly at different pH conditions (Supplementary Figure S7). Thus, these data indicate that the anionic Watson-Crick-like G•U^−^ can be a substantial driver of misincorporation errors during translation, transcription and possibly CRISPR/Cas off-target editing.

## CONCLUSION

In summary, like in B-DNA and A-RNA duplexes, tautomeric and anionic Watson-Crick-like G•T/U mismatches can form transiently and in low-abundance in RNA:DNA hybrids under solution conditions. Our study also uncovered a crucial distinction between these dynamics in B-DNA, A-RNA, and their hybrids. Specifically, anionic Watson-Crick-like G•U^−^ forms with a ten-fold higher propensity relative to G•T^−^ due to the greater ease of deprotonating U versus T. This discovery raises an intriguing prospect – the potential for anionic Watson-Crick-like G•U^−^ to play a more substantial role driving errors during translation, transcription, and CRISPR/Cas editing. This phenomenon may also contribute to the emergence of strand-specific disparities in the frequency of misincorporating dG•rU versus dT•rG during transcription. Noteworthy in this context is the observation that natural chemical modifications such as 5-oxyacetic acid uridine (cmo^5^ U), which recode translation, achieve this by modulating the propensities to form anionic Watson-Crick-like G•U^−^.(75) The high propensity to deprotonate U-N3 could explain the prevalence of anionic inverted wobble G•U mismatches in crystal structures of RNA.(78-80) Dissecting the role of these nucleic acid dynamics quantitatively will require examining whether altering these dynamics by changing sequence context, pH, and introducing chemical modifications results in predictable and corresponding changes in the kinetics of misincorporation and accommodation as done in prior studies of misincorporation during DNA replication.(8,32-34,45)

## DATA AVAILABILITY

A public git repository is available at https://github.com/alhashimilab/RNA-DNA-Hybrids.git (permanent DOI: 10.5281/zenodo.8278146) containing all source code and a minimum dataset sufficient to reproduce Figures 2-4, S1-S7 and all supplementary tables and statistics in this manuscript. Full data sets are available upon request.

## SUPPLEMENTARY DATA

Supplementary Figures S1-S7 and Supplementary Tables S1-S3 are available online.

## FUNDING

National Institutes of Health [R01GM089846 to H.M.A].

## Conflict of interest statement

None declared.

## ACKNOWLEDGEMENTS

We thank members of the Al-Hashimi laboratory for their assistance and critical comments on the manuscript. We would also like to thank Pablo Szekely (Duke University) for helpful discussions and Philip Benfey (Duke University) for his support and assistance. We would like to thank Ron Venters and the Duke Magnetic Resonance Spectroscopy Center for nuclear magnetic resonance resources. H.M.A. is a member of the New York Structural Biology Center. The data collected at NYSBC was made possible by a grant from ORIP/NIH facility improvement grant CO6RR015495. The NYSBC 700 MHz spectrometer was purchased with funds from NIH grant S10OD018509. O.S. is an Awardee of the Women’s Postdoctoral Career Development Award from the Weizmann Institute of Science.

## Notes

https://github.com/alhashimilab/RNA-DNA-Hybrids.git

### Competing Interest Statement

The authors have declared no competing interest.

