## Supplemental Figures and Tables for "NMR measurements of transient low-populated tautomeric and anionic Watson-Crick-like G·T/U in RNA:DNA hybrids: Implications for the fidelity of transcription and CRISPR/Cas9 gene editing"

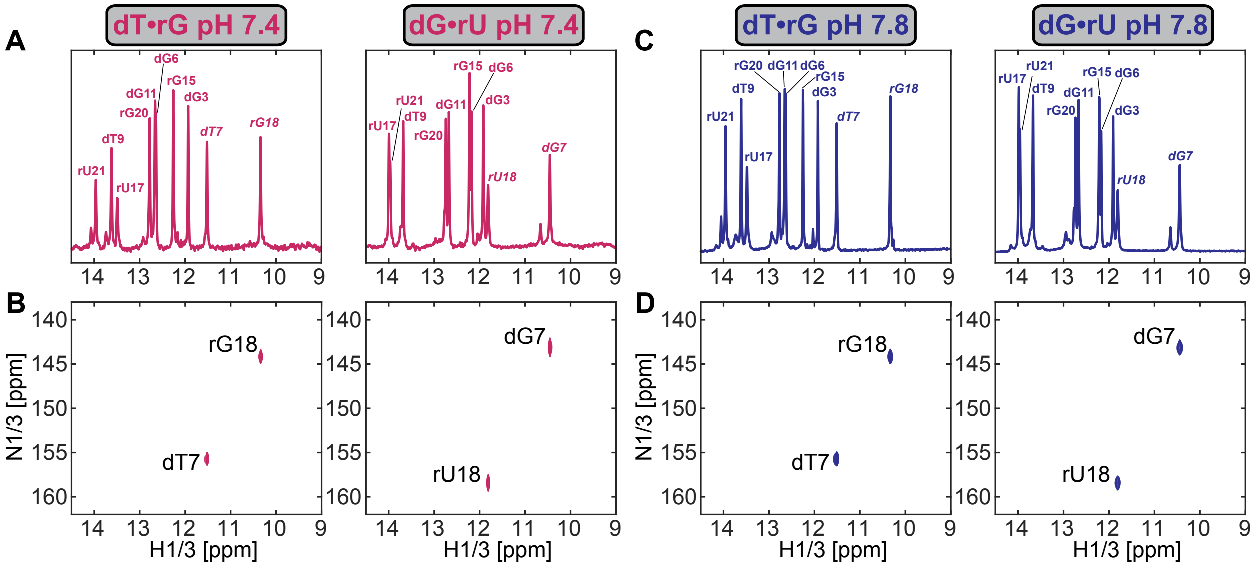


**Figure S1. NMR spectra of the two RNA:DNA hybrids.** (A) 1D $^{1}$H spectra and (B) 2D [$^{15}$N, $^{1}$H] HSQC spectra of the imino region of dT•rG (left) and dG•rU (right) measured at pH 7.4. Shown in the 2D HSQC spectra are the imino resonances of G-N1/H1 and T/U-N3/H3 targeted for relaxation dispersion measurements. The chemical shift of both G-H1 and T/U-H3 is up-field shifted relative to other resonances, indicative of the G•T/U wobble geometry. (C) 1D $^{1}$H spectra and (D) 2D [$^{15}$N, $^{1}$H] HSQC spectra of the imino region of dT•rG (left) and dG•rU (right) measured at pH 7.8.

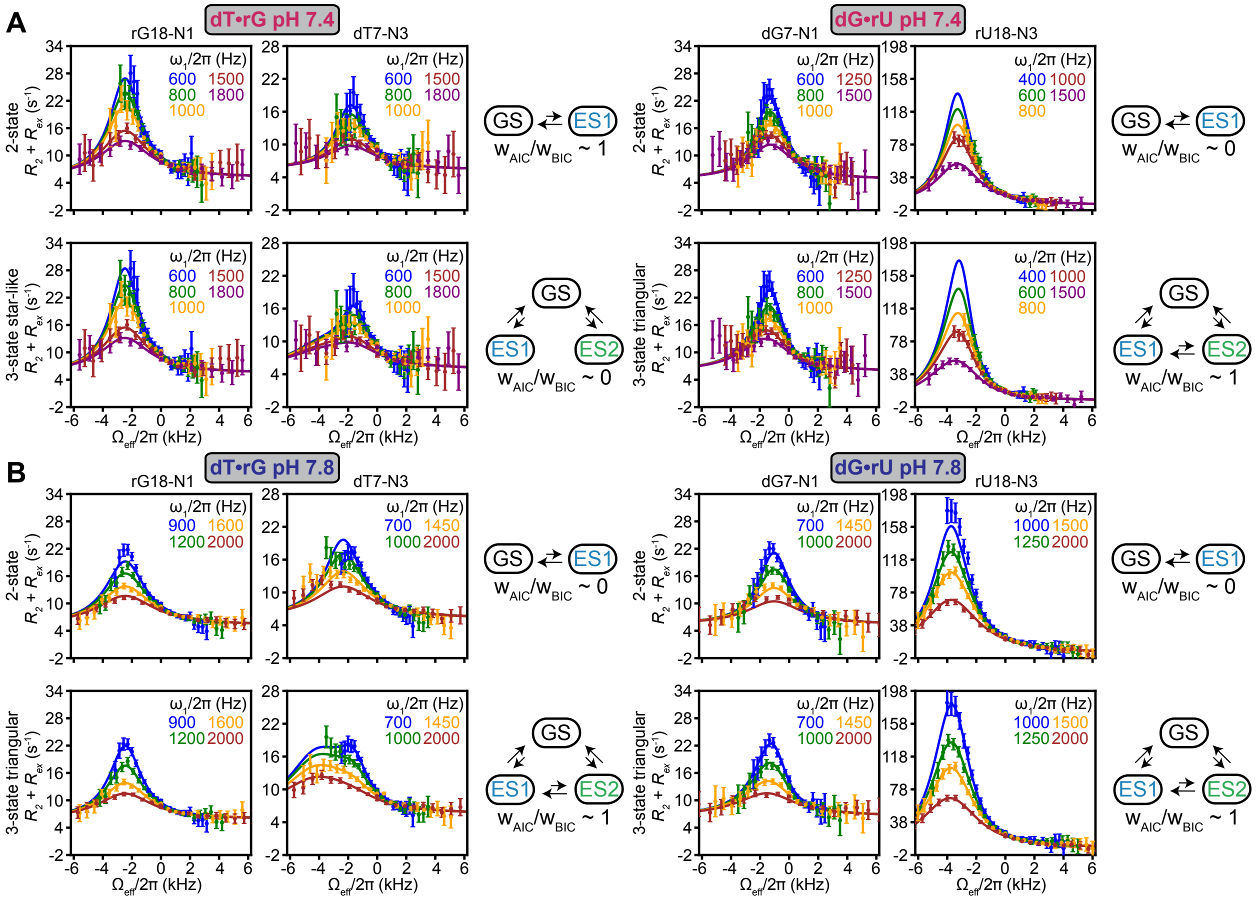

**Figure S2.** Comparison of global fits (solid lines) of the $^{15}$N $R_{1\rho}$ RD data (points) to 2- and 3-state exchange models. Spin-lock powers are color coded and error bars were obtained by propagating the error in $R_{1\rho}$ as described in Methods. Shown on the right of each comparison are the exchange topologies used in the fit along with the AIC and BIC weights ($w_{AIC}$ and $w_{BIC}$)(1,2) for each exchange model. The comparison is done for the dT•rG (left) and dG•rU (right) at pH 7.4 (A) and pH 7.8 (B). Based on the AIC and BIC statistical weights, a 2-state model was used to fit the $R_{1\rho}$ profiles of dT•rG at pH 7.4, whereas a 3-state triangular model was required to fit the $R_{1\rho}$ profiles for dG•rU at pH 7.4 and pH 7.8 and for dT•rG at pH 7.8.

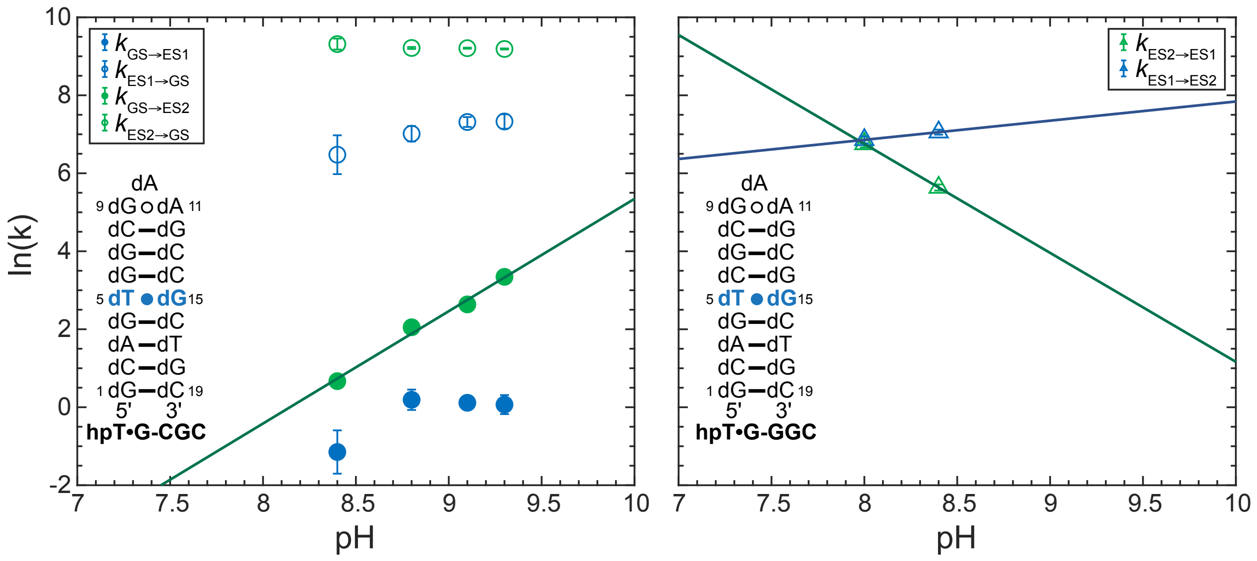


**Figure S3. Extrapolating exchange rates using pH dependent data.** Exchange rates were extracted from fits to off resonance $R_{1\rho}$ profiles measured in a prior study.(3) The natural logarithm of the exchange rates is plotted against the pH for hpTG-CGC (left) or hpTG-GGC (right). Forward and backward rates for ES1 (blue) and ES2 (green) are shown as filled and open symbols, respectively. Minor exchange rates between ES1 and ES2 are shown as open triangles. Solid lines are linear fits to the pH dependent data points.

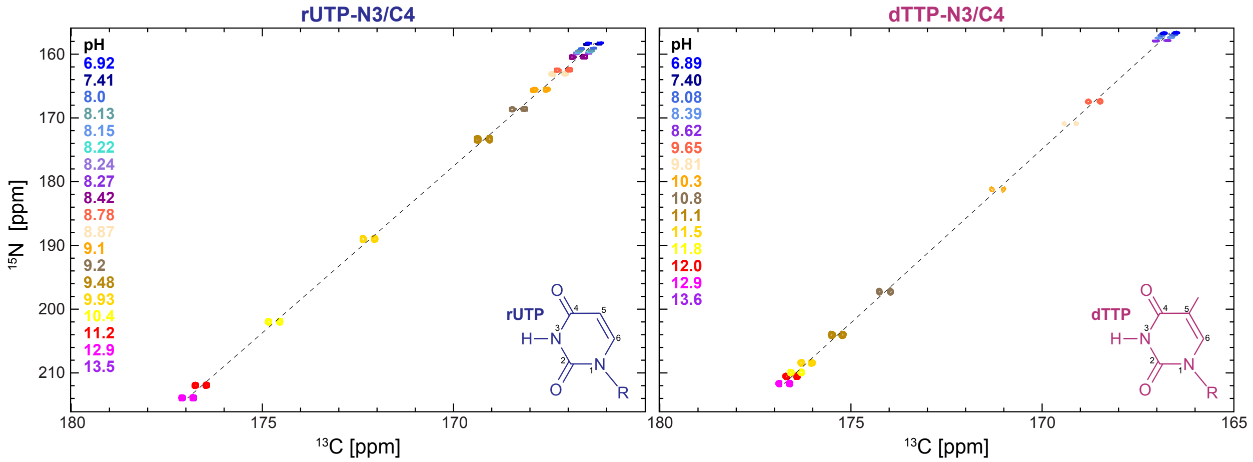


**Figure S4. NMR chemical shift perturbations used to measure the $pK_a$ of dTTP and rUTP.** NMR titrations of [$^{13}$C,$^{15}$N]-labeled rUTP (left) and dTTP (right), following the N3/C4 resonance in a 2D $^{13}$C-$^{15}$N CON HMQC spectrum. The sample pH values are color coded and indicated in the figure. Also shown is the chemical structure of the NTP (R = triphosphate).

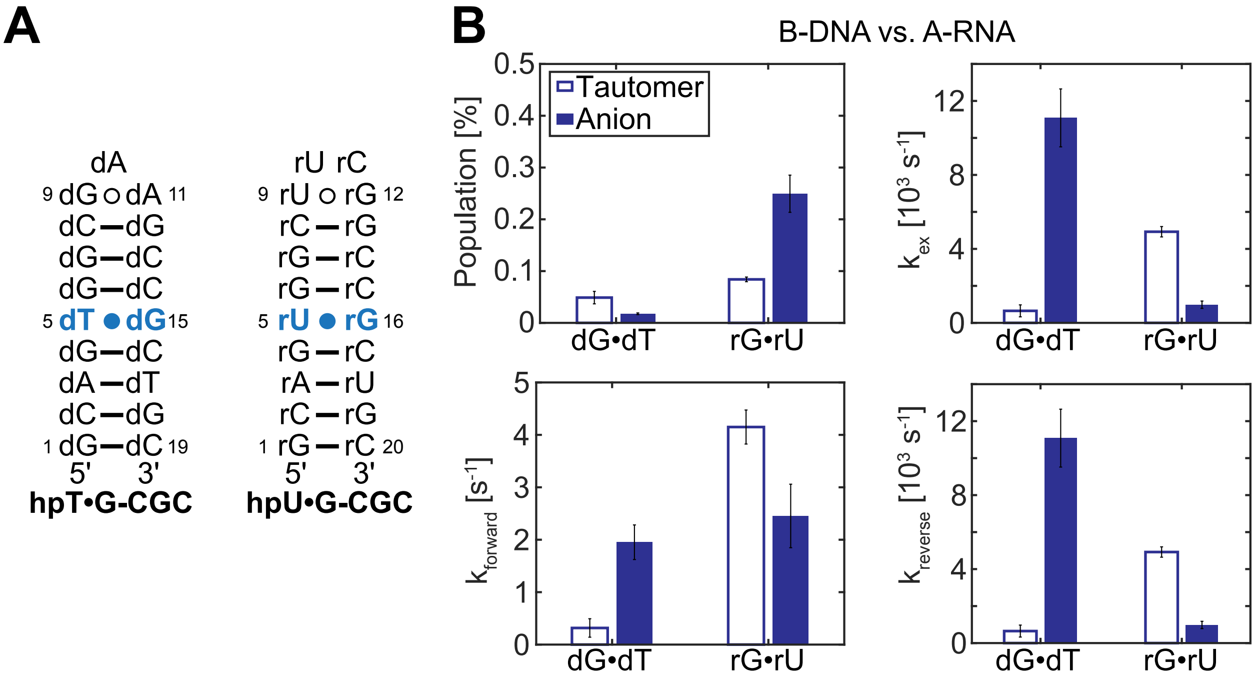


**Figure S5. Comparing propensities to form anionic G•T/U⁻ in A-RNA and B-DNA.** (A) B-DNA (left) and A-RNA (right) duplexes with a CGC trinucleotide sequence context for which the Watson-Crick-like dynamics of the G•T/U mismatches (in light blue) were reported previously.(3) (B) Comparison of exchange parameters reported in the prior study(3) from $R_{1\rho}$ measurements on B-DNA and A-RNA with the CGC sequence context at pH 8.4 and 10 °C. The populations and exchange rates are shown for the tautomeric (open bars) and anionic (full bars) states of dG•dT in B-DNA and rG•rU in A-RNA.

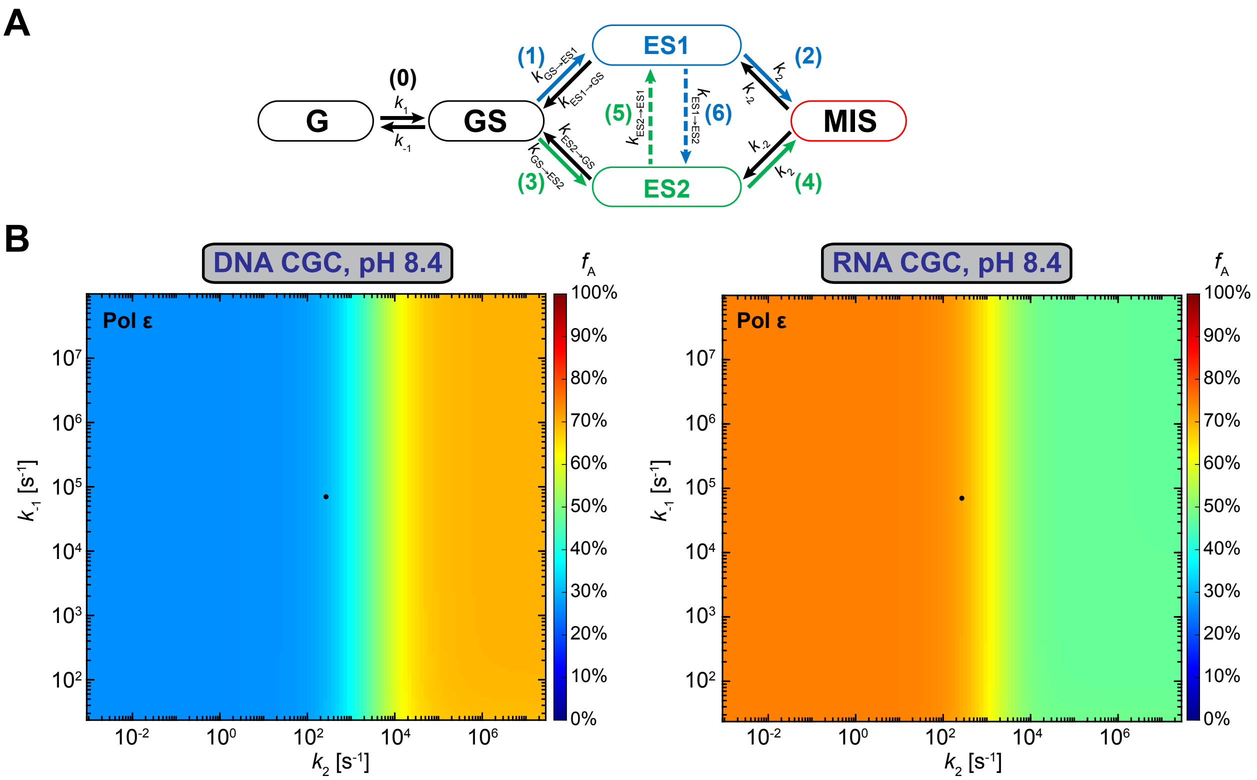


**Figure S6. Anionic Watson-Crick-like G•U⁻ contribution to misincorporation errors does not depend on initial binding step.** (A) General scheme for the kinetic pathways for misincorporating a G•T/U mismatch via tautomeric (in blue, ES1) and anionic (in green, ES2) Watson-Crick-like G•T/U conformational states. Shown are the binding step ($k_1$) to form the G•T/U wobble ground-state (GS); transition of the wobble into the tautomeric ($k_{GS\text{-}ES1}$) and anionic ($k_{GS\text{-}ES2}$) Watson-Crick-like G•T/U; and the kinetic step acting on the Watson-Crick-like species, which in DNA replication represents conformational changes in the DNA polymerase to form the catalytically active closed conformation. (B) Heat maps for the relative flux quantifying the fractional percentage ($f_A$) of misincorporations proceeding through the anionic Watson-Crick-like G•U⁻ conformation as a function of varying $k_{-1}$ and $k_2$ computed using the kinetic model presented in (A). Results are shown when using the chemical dynamics measured by NMR for dG•dT (DNA, left) and rG•rU (RNA, right) with a CGC trinucleotide sequence context(3) at pH 8.4. All other kinetic rate constants were assumed to be equal to those of DNA Polymerase ε.(4,5) Full circles indicate $k_{-1}$ and $k_2$ values for Pol ε.(4,5)

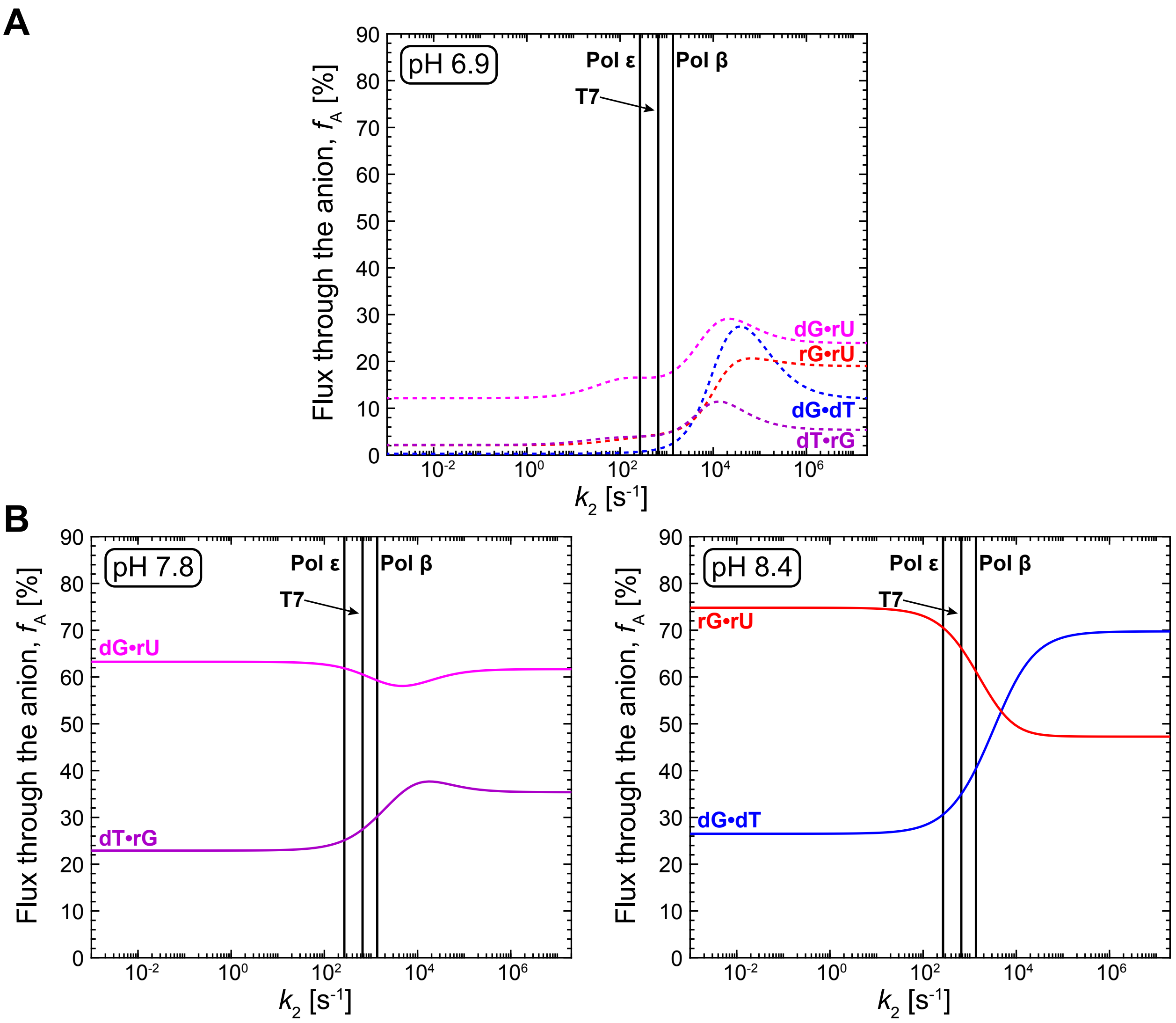


**Figure S7. Anionic Watson-Crick-like G•U⁻ contribution to misincorporation errors at different pH.** Relative flux quantifying the fractional percentage ($f_A$) of misincorporations proceeding through the anionic Watson-Crick-like G•U⁻ conformation as a function of varying $k_2$ computed using the kinetic model presented in Figure 4A. Results are shown when using the chemical dynamics measured by NMR for dG•dT (DNA) and rG•rU (RNA) with a CGC trinucleotide sequence context(3) and dG•rU and dT•rG in the two RNA:DNA hybrids examined in this work. The flux through the two pathways is computed using extrapolated exchange rates (dotted lines, see Materials and Methods) at pH 6.9 (A) or NMR-derived exchange rates (full lines) at pH 7.8 for the two hybrids (B, left), or at pH 8.4 for B-DNA and A-RNA (B, right). The calculations assume that the Watson-Crick-like G•T/U is an obligatory intermediate during misincorporation. Vertical lines indicate $k_2$ values for Pol ε,(4,5) T7,(6,7) and Pol β.(8)

**Table S1.** List of spin-lock powers ($\omega_1/2\pi$ in Hz) and offsets ($\Omega_{eff}/2\pi$ in Hz) used in off-resonance $^{15}$N $R_{1\rho}$ experiments.

| Nucleus | $\omega_1/2\pi$ (Hz), [$\Omega_{eff}/2\pi$ (Hz)] |
|---|---|
| dT•rG<br>rG18-N1<br>pH 7.4, 25 °C | 600, [-2097.0, -1864.0, -1631.0, -1398.0, -1165.0, -932.0, -699.0, -466.0, -233.0, -10.0, 10.0, 233.0, 466.0, 699.0, 932.0, 1165.0, 1398.0, 1631.0, 1864.0, 2097.0}<br><br>[800], {-2799.0, -2488.0, -2177.0, -1866.0, -1555.0, -1244.0, -933.0, -622.0, -311.0, -10.0, 10.0, 311.0, 622.0, 933.0, 1244.0, 1555.0, 1866.0, 2177.0, 2488.0, 2799.0]<br><br>1000, [-3501.0, -3112.0, -2723.0, -2334.0, -1945.0, -1556.0, -1167.0, -778.0, -389.0, -10.0, 10.0, 389.0, 778.0, 1167.0, 1556.0, 1945.0, 2334.0, 2723.0, 3112.0, 3501.0]<br><br>1500, [-5247.0, -4664.0, -4081.0, -3498.0, -2915.0, -2332.0, -1749.0, -1166.0, -583.0, -10.0, 10.0, 583.0, 1166.0, 1749.0, 2332.0, 2915.0, 3498.0, 4081.0, 4664.0, 5247.0]<br><br>1800, [-5600.0, -4900.0, -4200.0, -3500.0, -2800.0, -2100.0, -1400.0, -700.0, -10.0, 10.0, 700.0, 1400.0, 2100.0, 2800.0, 3500.0, 4200.0, 4900.0, 5600.0] |
| dT•rG<br>dT7-N3<br>pH 7.4, 25 °C | 600, [-2097.0, -1864.0, -1631.0, -1398.0, -1165.0, -932.0, -699.0, -466.0, -233.0, -10.0, 10.0, 233.0, 466.0, 699.0, 932.0, 1165.0, 1398.0, 1631.0, 1864.0, 2097.0]<br><br>800, [-2799.0, -2488.0, -2177.0, -1866.0, -1555.0, -1244.0, -933.0, -622.0, -311.0, -10.0, 10.0, 311.0, 622.0, 933.0, 1244.0, 1555.0, 1866.0, 2177.0, 2488.0, 2799.0]<br><br>1000, [-3501.0, -3112.0, -2723.0, -2334.0, -1945.0, -1556.0, -1167.0, -778.0, -389.0, -10.0, 10.0, 389.0, 778.0, 1167.0, 1556.0, 1945.0, 2334.0, 2723.0, 3112.0, 3501.0] |

| | |
|---|---|
| | 1500, [-5247.0, -4664.0, -4081.0, -3498.0, -2915.0, -2332.0, -1749.0, -1166.0, -583.0, -10.0, 10.0, 583.0, 1166.0, 1749.0, 2332.0, 2915.0, 3498.0, 4081.0, 4664.0, 5247.0]<br><br>1800, [-5600.0, -4900.0, -4200.0, -3500.0, -2800.0, -2100.0, -1400.0, -700.0, -10.0, 10.0, 700.0, 1400.0, 2100.0, 2800.0, 3500.0, 4200.0, 4900.0, 5600.0] |
| dG•rU<br><br>dG7-N1<br><br>pH 7.4, 25 °C | 600, [-2101.0, -1910.0, -1719.0, -1528.0, -1337.0, -1146.0, -955.0, -764.0, -573.0, -382.0, -191.0, -10.0, 10.0, 191.0, 382.0, 573.0, 764.0, 955.0, 1146.0, 1337.0, 1528.0, 1719.0, 1910.0, 2101.0]<br><br>800, [-2805.0, -2550.0, -2295.0, -2040.0, -1785.0, -1530.0, -1275.0, -1020.0, -765.0, -510.0, -255.0, -10.0, 10.0, 255.0, 510.0, 765.0, 1020.0, 1275.0, 1530.0, 1785.0, 2040.0, 2295.0, 2550.0, 2805.0]<br><br>1000, [-3498.0, -3180.0, -2862.0, -2544.0, -2226.0, -1908.0, -1590.0, -1272.0, -954.0, -636.0, -318.0, -10.0, 10.0, 318.0, 636.0, 954.0, 1272.0, 1590.0, 1908.0, 2226.0, 2544.0, 2862.0, 3180.0, 3498.0]<br><br>1250, [-4378.0, -3980.0, -3582.0, -3184.0, -2786.0, -2388.0, -1990.0, -1592.0, -1194.0, -796.0, -398.0, -10.0, 10.0, 398.0, 796.0, 1194.0, 1592.0, 1990.0, 2388.0, 2786.0, 3184.0, 3582.0, 3980.0, 4378.0]<br><br>1500, [-5247.0, -4770.0, -4293.0, -3816.0, -3339.0, -2862.0, -2385.0, -1908.0, -1431.0, -954.0, -477.0, -10.0, 10.0, 477.0, 954.0, 1431.0, 1908.0, 2385.0, 2862.0, 3339.0, 3816.0, 4293.0, 4770.0, 5247.0] |
| dG•rU<br><br>rU18-N3<br><br>pH 7.4, 25 °C | 400, [-1397.0, -1270.0, -1143.0, -1016.0, -889.0, -762.0, -635.0, -508.0, -381.0, -254.0, -127.0, -10.0, 10.0, 127.0, 254.0, 381.0, 508.0, 635.0, 762.0, 889.0, 1016.0, 1143.0, 1270.0, 1397.0]<br><br>600, -2101.0, -1910.0, -1719.0, -1528.0, -1337.0, -1146.0, -955.0, -764.0, -573.0, -382.0, -191.0, -10.0, 10.0, 191.0, 382.0, 573.0, 764.0, 955.0, 1146.0, 1337.0, 1528.0, 1719.0, 1910.0, 2101.0 |

| | |
|---|---|
| | 800, -2805.0, -2550.0, -2295.0, -2040.0, -1785.0, -1530.0, -1275.0, -1020.0, -765.0, -510.0, -255.0, -10.0, 10.0, 255.0, 510.0, 765.0, 1020.0, 1275.0, 1530.0, 1785.0, 2040.0, 2295.0, 2550.0, 2805.0<br><br>1000, [-3498.0, -3180.0, -2862.0, -2544.0, -2226.0, -1908.0, -1590.0, -1272.0, -954.0, -636.0, -318.0, -10.0, 10.0, 318.0, 636.0, 954.0, 1272.0, 1590.0, 1908.0, 2226.0, 2544.0, 2862.0, 3180.0, 3498.0]<br><br>1500, [-5247.0, -4770.0, -4293.0, -3816.0, -3339.0, -2862.0, -2385.0, -1908.0, -1431.0, -954.0, -477.0, -10.0, 10.0, 477.0, 954.0, 1431.0, 1908.0, 2385.0, 2862.0, 3339.0, 3816.0, 4293.0, 4770.0, 5247.0] |
| dT•rG<br><br>rG18-N1<br><br>pH 7.8, 25 °C | 900, [-3146.0, -2860.0, -2574.0, -2288.0, -2002.0, -1716.0, -1430.0, -1144.0, -858.0, -572.0, -286.0, -10.0, 10.0, 286.0, 572.0, 858.0, 1144.0, 1430.0, 1716.0, 2002.0, 2288.0, 2574.0, 2860.0, 3146.0]<br><br>1200, [-4202.0, -3820.0, -3438.0, -3056.0, -2674.0, -2292.0, -1910.0, -1528.0, -1146.0, -764.0, -382.0, -10.0, 10.0, 382.0, 764.0, 1146.0, 1528.0, 1910.0, 2292.0, 2674.0, 3056.0, 3438.0, 3820.0, 4202.0]<br><br>1600, [-5599.0, -5090.0, -4581.0, -4072.0, -3563.0, -3054.0, -2545.0, -2036.0, -1527.0, -1018.0, -509.0, -10.0, 10.0, 509.0, 1018.0, 1527.0, 2036.0, 2545.0, 3054.0, 3563.0, 4072.0, 4581.0, 5090.0, 5599.0]<br><br>2000, [-6996.0, -6360.0, -5724.0, -5088.0, -4452.0, -3816.0, -3180.0, -2544.0, -1908.0, -1272.0, -636.0, -10.0, 10.0, 636.0, 1272.0, 1908.0, 2544.0, 3180.0, 3816.0, 4452.0, 5088.0, 5724.0, 6360.0, 6996.0] |
| dT•rG<br><br>dT7-N3<br><br>pH 7.8, 25 °C | 700, [-2453.0, -2230.0, -2007.0, -1784.0, -1561.0, -1338.0, -1115.0, -892.0, -669.0, -446.0, -223.0, -10.0, 10.0, 223.0, 446.0, 669.0, 892.0, 1115.0, 1338.0, 1561.0, 1784.0, 2007.0, 2230.0, 2453.0] |

| | |
|---|---|
| | 1000, [-3498.0, -3180.0, -2862.0, -2544.0, -2226.0, -1908.0, -1590.0, -1272.0, -954.0, -636.0, -318.0, -10.0, 10.0, 318.0, 636.0, 954.0, 1272.0, 1590.0, 1908.0, 2226.0, 2544.0, 2862.0, 3180.0, 3498.0]<br>1450, [-5071.0, -4610.0, -4149.0, -3688.0, -3227.0, -2766.0, -2305.0, -1844.0, -1383.0, -922.0, -461.0, -10.0, 10.0, 461.0, 922.0, 1383.0, 1844.0, 2305.0, 2766.0, 3227.0, 3688.0, 4149.0, 4610.0, 5071.0]<br>2000, [-6996.0, -6360.0, -5724.0, -5088.0, -4452.0, -3816.0, -3180.0, -2544.0, -1908.0, -1272.0, -636.0, -10.0, 10.0, 636.0, 1272.0, 1908.0, 2544.0, 3180.0, 3816.0, 4452.0, 5088.0, 5724.0, 6360.0, 6996.0] |
| dG•rU<br>dG7-N1<br>pH 7.8, 25 °C | 700, [-2448.0, -2176.0, -1904.0, -1632.0, -1360.0, -1088.0, -816.0, -544.0, -272.0, -10.0, 10.0, 272.0, 544.0, 816.0, 1088.0, 1360.0, 1632.0, 1904.0, 2176.0, 2448.0]<br>1000, [-3501.0, -3112.0, -2723.0, -2334.0, -1945.0, -1556.0, -1167.0, -778.0, -389.0, -10.0, 10.0, 389.0, 778.0, 1167.0, 1556.0, 1945.0, 2334.0, 2723.0, 3112.0, 3501.0]<br>1450, [-5076.0, -4512.0, -3948.0, -3384.0, -2820.0, -2256.0, -1692.0, -1128.0, -564.0, -10.0, 10.0, 564.0, 1128.0, 1692.0, 2256.0, 2820.0, 3384.0, 3948.0, 4512.0, 5076.0]<br>2000, [-7002.0, -6224.0, -5446.0, -4668.0, -3890.0, -3112.0, -2334.0, -1556.0, -778.0, -10.0, 10.0, 778.0, 1556.0, 2334.0, 3112.0, 3890.0, 4668.0, 5446.0, 6224.0, 7002.0] |
| dG•rU<br>rU18-N3<br>pH 7.8, 25 °C | 1000, [-4004.0, -3718.0, -3432.0, -3146.0, -2860.0, -2574.0, -2288.0, -2002.0, -1716.0, -1430.0, -1144.0, -858.0, -572.0, -286.0, -10.0, 10.0, 286.0, 572.0, 858.0, 1144.0, 1430.0, 1716.0, 2002.0, 2288.0, 2574.0, 2860.0, 3146.0, 3432.0, 3718.0, 4004.0]<br>1250, [-4998.0, -4641.0, -4284.0, -3927.0, -3570.0, -3213.0, -2856.0, -2499.0, -2142.0, -1785.0, -1428.0, -1071.0, -714.0, -357.0, -10.0, 10.0, 357.0, 714.0, |

| | |
|---|---|
| | 1071.0, 1428.0, 1785.0, 2142.0, 2499.0, 2856.0, 3213.0, 3570.0, 3927.0, 4284.0, 4641.0, 4998.0]<br><br>1500, [-6006.0, -5577.0, -5148.0, -4719.0, -4290.0, -3861.0, -3432.0, -3003.0, -2574.0, -2145.0, -1716.0, -1287.0, -858.0, -429.0, -10.0, 10.0, 429.0, 858.0, 1287.0, 1716.0, 2145.0, 2574.0, 3003.0, 3432.0, 3861.0, 4290.0, 4719.0, 5148.0, 5577.0, 6006.0]<br><br>2000, [-7994.0, -7423.0, -6852.0, -6281.0, -5710.0, -5139.0, -4568.0, -3997.0, -3426.0, -2855.0, -2284.0, -1713.0, -1142.0, -571.0, -10.0, 10.0, 571.0, 1142.0, 1713.0, 2284.0, 2855.0, 3426.0, 3997.0, 4568.0, 5139.0, 5710.0, 6281.0, 6852.0, 7423.0, 7994.0] |

**Table S2.** Exchange parameters obtained from fitting $^{15}N$ $R_{1\rho}$ data in dG•rU and dT•rG at pH 7.4, 25 °C. GS corresponds to the wobble ground state of G•T/U, ES1 corresponds to the tautomeric Watson-Crick-like species which consists of $G^{enol}$•T/U and G•$T^{enol}$/$U^{enol}$ conformations in rapid equilibrium, and ES2 corresponds to the anionic G•$T^{-}$/$U^{-}$ conformation. Red $\chi^2$ is the reduced $\chi^2$ obtained on fitting the data. The exchange parameters that best fit the data based on statistical tests and chemical shift criteria as outlined in Methods are highlighted in bold.

| | Parameter | **dG•rU pH 7.4** | | **dT•rG pH 7.4** | |
|---|---|---|---|---|---|
| | | dG7-N1 | rU18-N3 | rG18-N1 | dT7-N3 |
| 2-state Individual Fitting | $p_{ES1}$ (%) | 0.201 ± 0.008 | 0.272 ± 0.009 | 0.082 ± 0.003 | 0.089 ± 0.007 |
| | $k_{ex,GS:ES1}$ ($s^{-1}$) | 6200 ± 400 | 5200 ± 300 | 6900 ± 600 | 8000 ± 1000 |
| | $\Delta\omega_{ES1}$ (ppm) | 19.4 ± 0.4 | 45.3 ± 0.4 | 35.0 ± 0.7 | 25 ± 1 |
| | $R_1$ ($s^{-1}$) | 2.65 ± 0.05 | 2.27 ± 0.07 | 2.18 ± 0.05 | 2.38 ± 0.06 |
| | $R_2$ ($s^{-1}$) | 5.4 ± 0.3 | 4.8 ± 0.4 | 5.3 ± 0.3 | 5.0 ± 0.4 |
| | Reduced $\chi^2$ | 0.56 | 0.61 | 0.55 | 0.61 |
| 2-state Shared Fitting | $p_{ES1}$ (%) | 0.241 ± 0.004 | | **0.083 ± 0.002** | |
| | $k_{ex,GS:ES1}$ ($s^{-1}$) | 6500 ± 200 | | **7200 ± 500** | |
| | $\Delta\omega_{ES1}$ (ppm) | 17.9 ± 0.3 | 46.1 ± 0.5 | **34.9 ± 0.7** | **25.7 ± 0.8** |
| | $R_1$ ($s^{-1}$) | 2.76 ± 0.04 | 2.41 ± 0.07 | **2.20 ± 0.05** | **2.34 ± 0.05** |
| | $R_2$ ($s^{-1}$) | 4.7 ± 0.2 | 3.9 ± 0.4 | **5.1 ± 0.2** | **5.4 ± 0.2** |
| | Reduced $\chi^2$ | 0.67 | | **0.58** | |
| 3-state | $p_{ES1}$ (%) | **0.19 ± 0.06** | | 0.083 ± 0.003 | |

| | | | | | |
|---|---|---|---|---|---|
| Shared Fitting | $p_{ES2}$ (%) | **0.12 ± 0.06** | | 0.010 ± 0.006 | |
| | $k_{ex,GS:ES1}$ ($s^{-1}$) | **4000 ± 900** | | 6600 ± 500 | |
| | $k_{ex,GS:ES2}$ ($s^{-1}$) | **5000 ± 2000** | | 15000 ± 11000 | |
| | $k_{ex,ES2:ES2}$ ($s^{-1}$) | **13000 ± 6000** | | - | |
| | $\Delta\omega_{ES1}$ (ppm) | **21 ± 3** | **36 ± 4** | 35.0 ± 0.7 | 23 ± 2 |
| | $\Delta\omega_{ES2}$ (ppm) | **1 ± 7** | **58 ± 7** | -0.1 ± 100 | 55 ± 20 |
| | $R_1$ ($s^{-1}$) | **2.54 ± 0.05** | **2.12 ± 0.07** | 2.17 ± 0.05 | 2.33 ± 0.07 |
| | $R_2$ ($s^{-1}$) | **5.6 ± 0.6** | **5.5 ± 0.5** | 5.4 ± 0.2 | 4.9 ± 0.6 |
| | Reduced $\chi^2$ | **0.52** | | 0.57 | |

**Table S3.** Exchange parameters obtained from fitting $^{15}N$ $R_{1\rho}$ data in dG•rU and dT•rG at pH 7.8, 25 °C. GS corresponds to the wobble ground state of G•T/U, ES1 corresponds to the tautomeric Watson-Crick-like species which consists of $G^{enol}$•T/U and G•$T^{enol}$/$U^{enol}$ conformations in rapid equilibrium, and ES2 corresponds to the anionic G•$T^{-}$/$U^{-}$ conformation. Red $\chi^2$ is the reduced $\chi^2$ obtained on fitting the data. The exchange parameters that best fit the data based on statistical tests and chemical shift criteria as outlined in Methods are highlighted in bold.

| | Parameter | **dG•rU pH 7.8** | | **dT•rG pH 7.8** | |
|---|---|---|---|---|---|
| | | dG7-N1 | rU18-N3 | rG18-N1 | dT7-N3 |
| 2-state Individual Fitting | $p_{ES1}$ (%) | 0.223 ± 0.009 | 0.42 ± 0.01 | 0.078 ± 0.003 | 0.075 ± 0.006 |
| | $k_{ex,GS:ES1}$ ($s^{-1}$) | 6200 ± 400 | 5300 ± 200 | 6000 ± 500 | 12000 ± 1000 |
| | $\Delta\omega_{ES1}$ (ppm) | 18.5 ± 0.4 | 51.9 ± 0.3 | 34.30 ± 0.6 | 35 ± 1 |
| | $R_1$ ($s^{-1}$) | 3.19 ± 0.04 | 2.33 ± 0.07 | 2.41 ± 0.03 | 2.68 ± 0.05 |
| | $R_2$ ($s^{-1}$) | 6.5 ± 0.3 | 4.8 ± 0.3 | 5.8 ± 0.2 | 4.3 ± 0.4 |
| | Reduced $\chi^2$ | 0.62 | 0.43 | 1.29 | 1.89 |
| 2-state Shared Fitting | $p_{ES1}$ (%) | 0.355 ± 0.007 | | 0.073 ± 0.002 | |
| | $k_{ex,GS:ES1}$ ($s^{-1}$) | 6900 ± 300 | | 8700 ± 600 | |
| | $\Delta\omega_{ES1}$ (ppm) | 14.8 ± 0.3 | 52.7 ± 0.4 | 34.7 ± 0.7 | 33.0 ± 0.8 |
| | $R_1$ ($s^{-1}$) | 3.42 ± 0.05 | 2.5 ± 0.1 | 2.48 ± 0.04 | 2.60 ± 0.04 |
| | $R_2$ ($s^{-1}$) | 5.3 ± 0.2 | 3.5 ± 0.5 | 5.2 ± 0.2 | 5.2 ± 0.2 |
| | Reduced $\chi^2$ | 1.00 | | 1.83 | |
| 3-state | $p_{ES1}$ (%) | **0.19 ± 0.05** | | **0.074 ± 0.007** | |

| | | | | | |
|---|---|---|---|---|---|
| Shared Fitting | $p_{ES2}$ (%) | **0.33 ± 0.05** | | **0.022 ± 0.005** | |
| | $k_{ex,GS:ES1}$ ($s^{-1}$) | **3200 ± 700** | | **5000 ± 700** | |
| | $k_{ex,GS:ES2}$ ($s^{-1}$) | **4300 ± 500** | | **9000 ± 2000** | |
| | $k_{ex,ES2:ES2}$ ($s^{-1}$) | **9000 ± 2000** | | **6000 ± 4000** | |
| | $\Delta\omega_{ES1}$ (ppm) | **21 ± 2** | **38 ± 4** | **35 ± 1** | **20 ± 3** |
| | $\Delta\omega_{ES2}$ (ppm) | **3 ± 2** | **57 ± 2** | **9 ± 10** | **62 ± 5** |
| | $R_1$ ($s^{-1}$) | **3.14 ± 0.04** | **2.14 ± 0.07** | **2.40 ± 0.04** | **2.49 ± 0.05** |
| | $R_2$ ($s^{-1}$) | **6.5 ± 0.4** | **5.9 ± 0.4** | **5.9 ± 0.2** | **5.4 ± 0.3** |
| | Reduced $\chi^2$ | **0.42** | | **1.25** | |
